# Multi-omics integration uncovers host-microbiota crosstalk underlying sexual differentiation in the shortfin eel *Anguilla bicolor pacifica*

**DOI:** 10.1101/2025.07.03.663112

**Authors:** Guo-Jie Brandon-Mong, Tzong-Huei Lee, Yi-Lung Chen, Tsun-Hsien Hsiao, Guan-Lun Wu, Yi-Li Lai, Chieh-Yin Weng, Ronnie G Gicana, Yu-Ting Yen, Yi-Fan Xu, Tzi-Yuan Wang, Yin-Ru Chiang

**Affiliations:** Biodiversity Research Center, Academia Sinica, Taipei 115, Taiwan; Institute of Fisheries Science, National Taiwan University, 106 Taipei, Taiwan; Department of Agricultural Chemistry, National Taiwan University, Taipei 106, Taiwan; School of Medicine, National Tsing Hua University, Hsinchu 300, Taiwan; Institute of Molecular and Cellular Biology, National Tsing Hua University, Hsinchu 300, Taiwan

**Keywords:** *Anguilla*, *Deinococcus*, gut microbiota, host-microbe interactions, sexual differentiation, sex hormones, steroid metabolism

## Abstract

The global decline in anguillid eel populations has intensified interest in understanding their biology for conservation and aquaculture. While host-gut microbiota interactions are well-characterized in homeotherms, these relationships remain poorly understood in poikilotherms during sexual differentiation. We examined gut microbiota dynamics across developmental stages in the shortfin eel *Anguilla bicolor pacifica*, which exhibits early sexual differentiation and a relatively short life cycle. Glass eels were cultivated in controlled freshwater conditions for three years, with sampling at key stages: glass eel, elver, sex-undetermined eel, and sex-determined eel. Full-length 16S rRNA gene sequencing revealed significant compositional shifts during development, with higher bacterial richness in adults versus younger eels. Early stages were dominated by Pseudomonadota, while sex-determined adults showed increased Deinococcota abundance. Network analysis identified *Deinococcus*, *Sphingomonas*, and *Variovorax* as key genera in sex-determined eels, with positive correlations between anti-Müllerian hormone gene expression and these taxa. We isolated 66 gut bacterial strains capable of metabolizing sex hormones under microaerobic conditions. These isolates, representing 22 genera across four phyla, demonstrated diverse metabolic capabilities from partial oxidation to complete steroid mineralization. Multiple strains achieved complete estradiol degradation as single isolates—a rare metabolic capability of environmental microorganisms. Comparative genomic analysis revealed widespread steroid-metabolizing genes, with *Deinococcus* species showing previously unreported hormone degradation capabilities. Our multi-omics analysis demonstrates that gut microbiota composition and function are intimately linked to eel sexual development, suggesting bidirectional host-microbe interactions influencing reproductive physiology. These findings advance understanding of host-microbiota interactions in aquatic vertebrates and provide implications for eel aquaculture and conservation.

## Introduction

While most anguillids are marine species, the genus *Anguilla* (eels; **Figure 1**) is unique in performing long-distance migrations between fresh and marine waters. *Anguilla* populations have declined dramatically worldwide due to high market demand and overfishing (Aalto et al., 2016; Drouineau et al., 2018; Marini et al., 2021) [1–3]. For example, the European eel (*Anguilla anguilla*) has experienced a population decline exceeding 90% since the 1980s and has been classified as critically endangered on the International Union for Conservation of Nature (IUCN) red list since 2008 (Dekker, 2003; IUCN 2023) [4, 5]. Similarly, the Japanese eel (*Anguilla japonica*), a temperate Asian species, is listed as endangered, with its population decreasing by more than 50% over three decades (IUCN 2023) [5].

**Figure 1.**
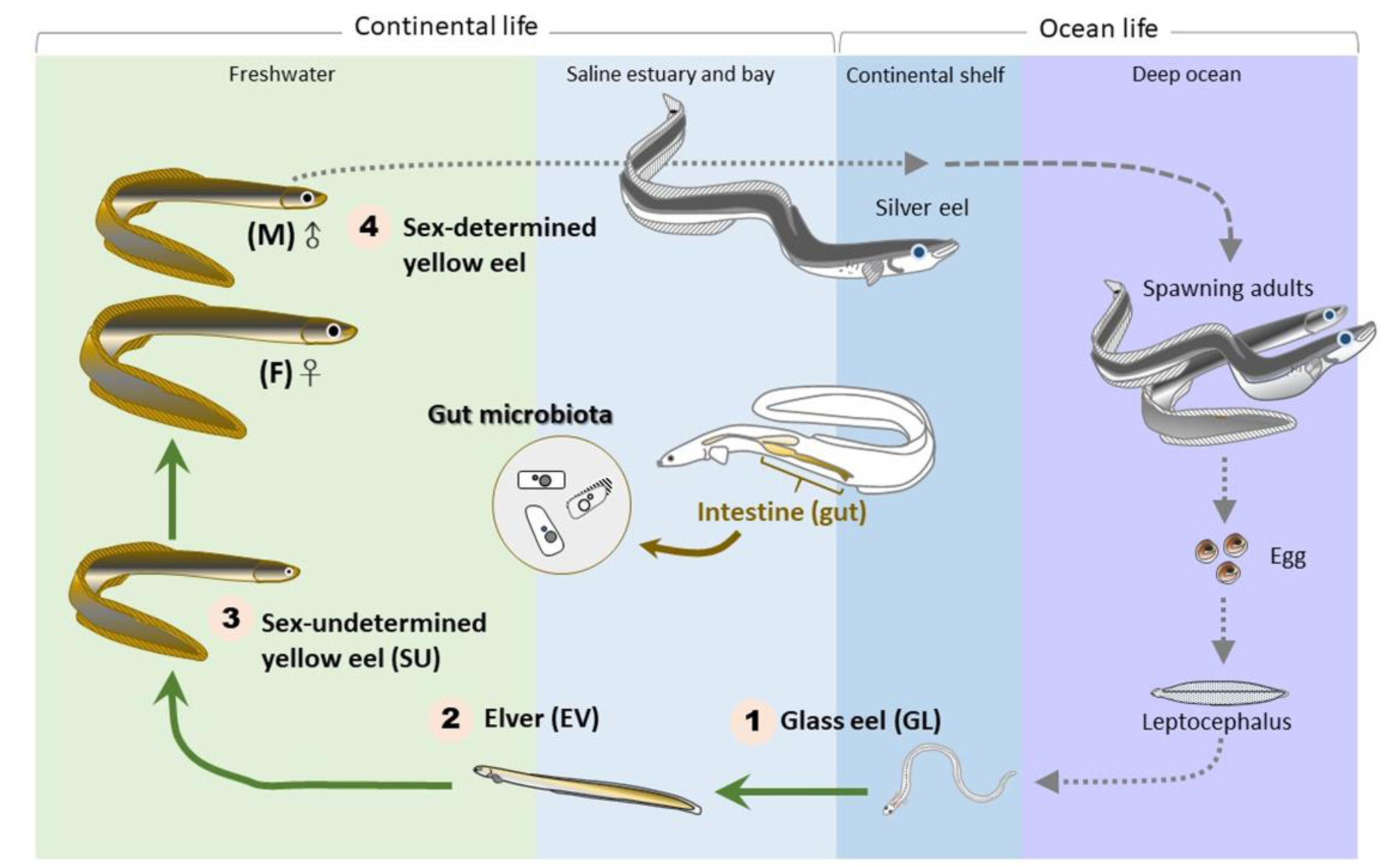
The complete life cycle of *Anguilla* spp. The numbers represent eel collected at different development stages: (1) GL, glass eel stage; (2) EV, elver stage; (3) SU, sex-undetermined juveniles; and (4) M and F, sex-determined male and female adults.

Unlike the extensively studied Japanese eel, the ecology and physiology of the tropical Asian shortfin eel (*Anguilla bicolor*) remain poorly understood. Currently listed as near threatened (IUCN 2023) [5], *A. bicolor* exhibits distinct life history traits compared to *A. japonica*. In general, *A. bicolor* undergoes sexual differentiation earlier than *A. japonica*. *A. bicolor* typically undergoes sex differentiation up to 4 years, while *A. japonica* typically undergoes sex differentiation up to 7 years, often requiring extended periods to achieve full differentiation (Arai & Chino, 2013; Chai & Arai, 2018; Horiuchi et al., 2022) [6–8]. Additionally, sexually differentiated *A. bicolor* are frequently encountered in freshwater environments (Arai & Abdul Kadir, 2017) [9]. Beyond their extraordinary migratory ecology, eels exhibit remarkably complex life cycles and sexual development patterns. Their reproductive biology is governed by sophisticated hormonal mechanisms that have been most extensively studied in Japanese eels. This research has revealed pronounced sexual dimorphism, with females typically achieving enhanced growth rates and larger body sizes compared to males (Kotake et al., 2007) [10]. These morphological differences are complemented by distinct sex-specific molecular markers: males demonstrate elevated expression of the gonadal anti-Müllerian hormone (*amh*) gene (Lin et al., 2021) [11], while females exhibit higher expression levels of gonadal vitellogenin receptor (*vtgr*), which encodes an oocyte membrane-specific receptor essential for egg development (Lai et al., 2022) [12].

The primary sex steroids in eels, particularly Japanese eels, include two androgens—testosterone and 11-ketotestosterone—and one estrogen, 17*β*-estradiol (hereafter referred to as estradiol) (Hwang et al., 2022) [13]. Testosterone functions both as a principal androgen and as a precursor hormone in eels, contributing to the development of secondary sexual characteristics (Peñaranda et al., 2014) [14]. Additionally, testosterone serves as a biochemical precursor that can be converted to other sex steroids, including 11-ketotestosterone and estradiol. 11-Ketotestosterone, derived from testosterone, is critical for testicular development and spermatogenesis, particularly during the final stages of sperm maturation (Baeza et al., 2015) [15]. Similar to mammals, estradiol serves as the primary female sex hormone in eels. Produced in the ovaries through aromatization of testosterone, this female-specific steroid is essential for ovarian development and vitellogenesis (yolk formation in eggs) (Palstra et al., 2022; Peñaranda et al., 2014) [14, 16]. It stimulates the liver to produce vitellogenin, a protein crucial for egg development. Using Japanese eel as a model organism, researchers have observed that concentrations of these sex steroids increase significantly during sexual maturation, regulating the silvering process—the metamorphosis from yellow to silver eels preceding their oceanic spawning migration (Aroua et al., 2005; Han et al., 2003; Sudo et al., 2012; Sudo & Yada et al., 2022) [17–20]. The balance between these hormones shifts dramatically throughout the eel’s unique life cycle, particularly during the silvering phase (transition from yellow to silver eel). This metamorphic transformation is characterized by significant digestive tract regression, with 11-ketotestosterone playing a key role in inducing gut degeneration alongside other silvering-related changes such as eye enlargement and gonadal development (Sudo et al., 2012) [19]. The progressive gut atrophy serves as an adaptive preparation for the extensive non-feeding migration, during which silver eels must survive solely on accumulated energy stores during their oceanic spawning journey.

Like all complex organisms, eels exist not as isolated biological entities but as holobionts— integrated systems of host and associated microbial communities (Simon et al., 2019) [21]. This host-microbiota framework offers a promising lens for understanding sexual differentiation and reproductive physiology in anguillid eels. Accumulating evidence demonstrates that gut microbiota significantly influence critical host functions, particularly reproduction and endocrine regulation (Fukui et al., 2018) [22]. Notably, recent studies have shown that gut microbiota can actively modulate host hormone profiles through enterohepatic circulation, creating bidirectional communication between microbial communities and the host endocrine system (Hsiao et al., 2023; Chen et al., 2024) [23, 24]. This regulatory capacity extends to fish sex development, where gut bacteria have been shown to play important roles (Li et al., 2021; Meng et al., 2022; Piazzon et al., 2019) [25–27].

Despite these advances, the relationship between host sex hormones and gut microbiota composition remains poorly understood in aquatic animals, particularly in anguillid eels. Current research on eel gut microbiota is limited to only six studies spanning different species and life stages: juvenile of *A. anguilla* (Bertucci et al., 2022) [28], juvenile through silver adult of *A. anguilla* (Huang et al., 2018) [29], juvenile of *A. australis* (Kusumawaty et al., 2023) [30], glass eel of *A. rostrata* (Liang et al., 2023) [31], immature and adult of *A. marmorata* (Lin et al., 2019) [32], and sex-determined adult of *A. japonica* (Zhu et al., 2021) [33]. Critically, research on *A. bicolor* gut microbiota beyond the juvenile (i.e., elver) stage remains absent (Kusumawaty et al., 2023) [30]. Further complicating our understanding, previous studies were conducted across varied environmental conditions—with factors such as pollutants, habitat conditions, salinity, and temperature all capable of influencing gut microbiota composition. This environmental heterogeneity has left many fundamental aspects of eel-microbiota interactions unexplored. Controlled environment studies could therefore provide crucial insights into the specific bacterial taxa and their functional roles throughout eel development.

Here, we hypothesize that host-gut microbiota interactions drive developmental programming and sexual differentiation in the shortfin eel *Anguilla bicolor pacifica*. We aimed to: (1) characterize the interactions between sex hormones and gut microbiota across different life stages, and (2) identify key gut microbes capable of hormone metabolism in sex-determined *A. bicolor* eels. We are particularly interested in understanding the dynamic changes in gut microbiota composition during sexual differentiation and elucidating the functional role of gut microbiota in modulating eel sexual development. To minimize confounding environmental factors that could affect wild populations, we sourced *A. bicolor* specimens from a controlled aquaculture facility. We analyzed gut microbial communities at each developmental stage using full-length 16S rRNA PacBio sequencing and the QIIME2-DADA2 pipeline. Additionally, we employed a culture-dependent approach to isolate hormone-metabolizing bacterial strains from adult eels. Our analysis revealed dynamic shifts in gut microbiota composition throughout development, with a remarkable increase in *Deinococcus* abundance in adults and a corresponding decrease in *Pseudomonas* over the developmental trajectory. From adult eels, we successfully collected 66 gut bacterial strains (spanning four phyla), all capable of metabolizing sex hormones. Functional genomics analysis revealed the prevalence of steroid-degrading genes in these microbes, supporting their role in hormone metabolism. Together, our multi-omics approach demonstrates that host-gut microbiota interactions play a crucial role in developmental programming and sexual differentiation of *A. bicolor*, providing valuable insights for both eel ecophysiology and sustainable aquaculture practices.

## Materials and methods

### Eel culture system

Glass eels (*A. bicolor*) were collected from an estuary in Hualien, eastern Taiwan, and transported to a fish farm with non-polluted stream water in Hsinchu, Taiwan. Day 0 was designated as the arrival date of glass eels at the farm. The cultivation process proceeded through three stages: 1) Glass eels (GL) were reared for 6 months in freshwater tanks (10-15 cm depth, 20-25°C); 2) Resulting elvers (EV, fingerlings) were maintained in 1-2 m depth tanks for 1 year; 3) sex-undetermined eels (SU) were grown in 2-3 m depth tanks for 2 years. Specimens were collected at four developmental stages: glass eels (GL; day 0), elvers (EV; 6 months), sex-undetermined eels (SU; 1.5 years), and sex-determined eels (females and males; 3 years). Glass eels were fed red worms, while later stages received commercial fish feed pellets.

### Ethics statement

All procedures (Protocol ID: 22-11-1925, 22-11-1926) were approved by the Institutional Animal Care & Use Committee of Academia Sinica, Taiwan.

### Sample collection

Forty-four *A. bicolor* eels at different developmental stages were sampled: glass eels (GL, n = 6), elvers (EV, n = 8), sex-undetermined eels (SU, n = 10), sex-determined males (M, n = 10), and sex-determined females (F, n = 10). Eels were euthanized via hypothermia using ice-slurry immersion, followed by morphological measurements (body weight and length). Blood samples were collected from the tail of sex-determined eels (M and F) in heparinized tubes, centrifuged (3000 rcf, 10 minutes), and the plasma was stored at -20°C for hormone quantification. Tissue samples were processed as follows: 1) Gonad tissue: preserved in RNAlater overnight at 4°C, then stored at -20°C for RNA extraction; 2) Proximal intestine: dissected to expose luminal surface, stored at -20°C for DNA extraction; 3) Distal intestine: maintained on ice for same-day bacterial culture.

### RT-qPCR analysis

Expression levels of anti-müllerian hormone (*amh*) and vitellogenin receptor (*vtgr*) genes were quantified using RT-qPCR. Total RNA was extracted from gonad tissue using Direct-zol RNA Kits (R2052, ZymoResearch) and purified with Turbo DNA-free Kit (Thermo Fisher Scientific). First-strand cDNA synthesis was performed using SuperScript® IV System with oligo-dT primers (Thermo Fisher Scientific). Gene expression was analyzed using Power SYBR® Green PCR Master Mix with qPCR primers [Forward (*vtgr*): 5’-GCTCATAGACCGCAAGACC-3’; Reverse (*vtgr*): 5’-GCCTTACACACGCCAGAAGT-3’ (Morini et al., 2020) [34]; Forward (*amh*): 5’-TCCTGGTCAGCACTGCGTATC-3’; Reverse (*amh*): 5’-TCCCGCACCGACAGACA-3’; Forward (*ef1a*): 5’-TGTGGGAGTCAACAAGATGGA-3’ (Lin et al., 2021) [11]; Reverse (*ef1a*): 5’-CTCAAAACGCTTCTGGCTGTA-3’ (Lin et al., 2021) [11]] in a QuantStudioTM 5 Real-Time PCR System (Thermo Fisher Scientific), with elongation factor 1α (*ef1a*) serving as the internal control.

### DNA extraction and 16S rDNA amplicon sequencing

Genomic DNA was extracted from five sample types (eel gut, freshwater, seawater, red worms, and fish feed pellets) using PowerFecal DNA isolation kit (Qiagen, Germany). The full-length 16S rRNA gene was amplified using universal primers with PacBio overhang adapters and sample-specific barcodes: Forward (27F): 5’-AGAGTTTGATCCTGGCTCAG-3’; Reverse (1492R): 5’-GGTTACCTTGTTACGACTT-3’ (Lane, 1991) [35]. PCR amplification was performed in duplicate using SuperRed PCR Master Mix (BIOTOOLS Co., Ltd., Taiwan) with the following touchdown protocol: Initial denaturation at 95 °C for 30 s; 16 cycles of 10 s at 95 °C, 30 s at 62 °C with 1 °C decreasing delta temperature every subsequent cycle (final temperature was 46 °C), 1 min at 72 °C; followed by 25 cycles of 10 s at 95 °C, 30 s at 46 °C, 1 min at 72 °C; and a final elongation step for 5 min at 72 °C. Successful amplification (∼1,500 bp) was confirmed by 1.0% agarose gel electrophoresis. Purified PCR products (AMPure PB Beads) were sequenced using PacBio technology (BIOTOOLS Co., Ltd.). 16S rRNA gene sequence data are available in the Sequence Read Archive (SRA) under BioProject ID PRJNA1244584, with accession numbers ranging from SRR32945574-SRR32945632.

### Eel species identification

To identify eel species, the cytochrome oxidase subunit 1 (*COI*) gene was PCR amplified from DNA samples using SuperRed PCR Master Mix and universal primers (Forward LCO1490: 5’-GGTCAACAAATCATAAAGATATTGG-3’ and Reverse HCO2198: 5’-TAAACTTCAGGGTGACCAAAAAATCA-3’) described by Folmer et al. (1994) [36]. The PCR followed a touchdown protocol: Initial denaturation at 95 °C for 30 s; 16 cycles of 10 s at 95 °C, 30 s at 62 °C with 1 °C decreasing delta temperature every subsequent cycle (final temperature was 46 °C), 1 min at 72 °C; followed by 40 cycles of 10 s at 95 °C, 30 s at 46 °C, 1 min at 72 °C; and a final elongation step for 5 min at 72 °C. Successful amplification (∼700 base pairs) was confirmed using 1.0% agarose gel electrophoresis. After the purification of the PCR products, Sanger sequencing was performed, and a maximum likelihood phylogenetic tree was constructed using MEGA software (Tamura et al., 2021) [37]. The analysis included NCBI *COI* reference sequences from seven eel species (*A. japonica* [LC782003.1, KU168670.1], *A. marmorata* [MZ435985.1, MW283183.1], *A. anguilla* [KU927487.1, KU927488.1], *A. bicolor pacifica* [LC548771.1, MT647227.1], *A. bicolor bicolor* [PP352530.1, MZ312365.1], and *A. rostrata* [MW822312.1, MW822295.1]), with *Conger* (MN711446.1) serving as an outgroup for comparison.

### Gut microbiota analysis

Raw PacBio reads were processed using QIIME2 pipeline (Hall & Beiko, 2018) [38]. Sequences were filtered to remove: 1) Those outside 1000-1600 bp length range; 2) Primers and chimeric sequences; 3) Non-bacterial sequences (chloroplast, eukaryote, mitochondria, unknown taxa). Filtered sequences were denoised using DADA2 to generate amplicon sequence variants (ASVs) (Callahan et al., 2017) [39]. Species-level taxonomic classification used bacterial 16S rRNA RefSeq sequences from NCBI nucleotide database, with taxonomic data retrieved via NCBI Datasets Command Line Tool. Taxonomic classification thresholds were: Species: ≥98.7%; Genus: ≥94.5%; Family: ≥86.5%; Order: ≥82.0%; Class: ≥78.5%; Phylum: ≥75.0% (Barco et al., 2020; Chun et al., 2018; Yarza et al., 2014) [40–42]. ASVs were binned into taxonomic count tables using Mothur phylotype command (Schloss et al., 2009) [43]. Rarefaction curve analysis was conducted to validate sequencing depth adequacy for gut microbiota analysis. Samples were rarefied to 2,499 reads for diversity analyses, including Shannon’s diversity index (alpha diversity) and Bray-Curtis dissimilarity (beta diversity). The LEfSe-based analysis (lefser) was conducted to identify biomarkers (e.g., gut microbial genera) that statistically differentiate sex-determined eels (M & F) and younger eels (GL, EV & SU) (Khleborodova et al., 2024; Segata et al., 2011) [44, 45]. Within lefser, a non-parametric Kruskal-Wallis test was conducted to detect gut bacterial genera with significant differential abundance between the two groups (Khleborodova et al., 2024) [44]. Finally, Linear Discriminant Analysis (LDA) was performed in lefser to estimate the effect size of each differentially abundant genus and identify biomarkers with large differences between groups (Khleborodova et al., 2024) [44]. Venn diagram analysis was conducted to identify core bacterial genera persistently present across all developmental stages.

### Correlation analysis between gene expression and gut microbiota

Two correlation analyses were performed: 1) Among gut bacterial genera across developmental stages; 2) Between sex-specific gene expression (*amh* & *vtgr*) and gut bacterial genera across eel life stages (GL, EV, SU, M, F). Correlations were calculated using FastSpar (based on SparCC) with 1,000 bootstrap replicates (Friedman et al., 2012; Watts et al., 2019) [46, 47]. Significant bacterial interactions (*p* < 0.05) were used to calculate network centrality scores (degree, closeness, betweenness, and eigenvector) using Gephi (Bastian et al., 2009) [48]. For the network interpretations, each node represents a microbial genus, with node color and size indicating the eigenvector centrality score (red: higher score, blue: lower score). The edge color represents the interaction strength between nodes, reflecting correlations between microbial genera. Correlations between microbial genera and the expression of *amh* and *vtgr* are indicated as correlation coefficients (r values). Correlation coefficients (r) were interpreted as weak (0.1-0.3), moderate (0.3-0.7), or strong (0.7-1.0). Asterisks indicate significance levels for correlations between microbial genera and the expression of *amh* and *vtgr*: \**p* < 0.05; \*\**p* < 0.01; \*\*\**p* < 0.001.

### Culture-dependent approaches for isolation of hormone-metabolizing bacteria

There were three sequential strategies for isolating steroid-metabolizing microbes. First, intestinal microbiota was cultivated under three physiological conditions: aerobic, anaerobic, and fermentative growth, with testosterone, 11-ketotestosterone or estradiol provided as the specific steroids in all conditions. Second, single bacterial colonies were isolated from Tryptic Soy Agar (TSA) plates to obtain pure cultures. Finally, the steroid-metabolizing activities of each isolate were validated using silica-gel 60 F254 thin-layer chromatography (TLC) (Merck). The details are as follows:

Eel intestines were dissected, placed in sterile Petri dishes, and longitudinally opened under aseptic conditions. The tissue was suspended in sterile saline (1:10 w/v), vortexed to release microbes, and allowed to settle for 1 min to remove debris. The supernatant was pooled for inoculation. Intestinal microbiota were inoculated under three conditions: (1) aerobic culture: 1 mL suspension was inoculated into the phosphate-buffered chemically defined medium (Chen et al., 2017; Hsiao et al., 2022) [49, 50]; (2) denitrifying culture: 1 mL suspension was inoculated into a denitrification medium (Wei et al., 2018) [51]; (3) anaerobic fermentation: 4 mL suspension was added to an anaerobic bottle, flushed with nitrogen for 30 min, and inoculated into DCB-1 medium (Hemme et al., 2011) [52]. All cultures were supplemented with 1 mM of testosterone, 11-ketotestosterone or estradiol and incubated at 30°C, 100 rpm, with sub-samples collected on days 0, 1, 3, 5, 7, 14 and 21.

Steroid of each inoculation was extracted via ethyl acetate and the concentrations were monitored using TLC plates. Once depletion was observed, serial dilutions (10¹–10⁵) were spread on TSA for isolation. Single colonies with distinct morphologies were subcultured and purified through quadrant-streaking method. Each isolate was inoculated in phosphate-buffered chemically defined medium (Chen et al., 2017; Hsiao et al., 2022) [49, 50] supplemented with 1 mM of testosterone, 11-ketotestosterone or estradiol at 30°C with shaking at 100 rpm. Sub-sampling was performed on days 0, 3, and 7, with culture extracts obtained using ethyl acetate. Steroid conversion by each isolate was verified by spotting the ethyl-acetate extracts onto TLC plates and comparing the disappearance of parent steroids and appearance of reference standards under UV illumination.

To confirm, verify and strengthen our discoveries, in total, 11 *Deinococcus* strains were purchased from the Bioresource Collection and Research Center (BCRC), Hsinchu, Taiwan, including *Deinococcus radiodurans* (BCRC num: 12827^T^; ATCC13939), *Deinococcus radiopugnans* (12856^T^; ATCC19172), *Deinococcus grandis* (17376^T^; ATCC43672), *Deinococcus proteolyticus* (17377^T^; MRP), *Deinococcus geothermalis* (17378^T^; DSM11300), *Deinococcus indicus* (17379^T^; DR1), *Deinococcus murrayi* (17380^T^; DSM11303), *Deinococcus deserti* (17541^T^; VCD115), *Deinococcus navajonensis* (17558^T^; CGMCC1.12543), *Deinococcus hohokamensis* (17559^T^; CGMCC4.7563), and *Deinococcus ficus* (17568^T^; CC-FR2-10). Link: https://catalog.bcrc.firdi.org.tw/KeywordResult?searchType=1&key=deinococcus&bno=&strainall=all.

Each strain was cultured in R2A medium and purified through quadrant streaking. Then, single colony was cultured in phosphate-buffered chemically defined medium (Chen et al., 2017; Hsiao et al., 2022) [49, 50] with 1% oxygen with either testosterone, 11-ketotestosterone or estradiol for 7 days. Steroid degradation was monitored using TLC.

Steroid-metabolizing isolates were identified through 16S rRNA gene sequencing. Genomic DNA was extracted using the Presto Mini gDNA Bacteria Kit, followed by PCR amplification of the 16S rRNA gene using primers 27F (5’-AGAGTTTGATCMTGGCTCAG-3’) and 1492R (5’-TACGGYTACCTTGTTACGACTT-3’) (Lane, 1991) [35]. The amplified products were sequenced using the Sanger method, and the resulting sequences were analyzed using BLASTn against the NCBI database for taxonomic identification.

### Nanopore whole-genome sequencing

To identify bacterial genes associated with sex hormone metabolism, 21 bacterial strains isolated from eel gut microbiota were selected for whole-genome sequencing. Genomic DNA was extracted from bacteria single isolates with testosterone/estradiol metabolism function. Subsequently, 0.3-1μg of genomic DNA were barcoded with Native Barcoding Kit (SQK-NBD104, SQK-NBD114), subjected to end-repair and ligation using the KAPA Hyper Prep Kit (Cat#KR0961, Kapa Biosystems, Wilmington, MA, USA) according to the supplier’s guidelines. The prepared DNA library was combined with LB and SQB buffer from the Nanopore Ligation Sequencing Kit (SQK-LSK109, Oxford Nanopore Technologies, UK), loaded into a flow cell (R9.4.1; FLO-MIN106), and sequenced using MinION devices over a period of 24-72 hours. Guppy was used with the High Accuracy (HAC) basecalling model for basecalling and demultiplex reads by identifying barcodes from the basecalled results, separating reads according to their barcode sequences. The raw Fastq files were quality-filtered: 1) the first 100 bp of both ends of all reads were trimmed to remove primer, barcodes and adapters; 2) filtlong was used to remove reads shorter than 1000 bp, reads with a mean quality score below 10 and reads with the lowest-quality 10% of the data by base content (Wick, 2017) [53]. Flye was used for genome assembly (Kolmogorov et al., 2019) [54]. Racon was performed 3 rounds of polishing by aligning the original long reads (raw ONT fastq file) back to the assembly (output file from Flye) and correcting errors based on these alignments (Vaser et al., 2017) [55]. Medaka was performed to generate a highly accurate final consensus from the polished assembly from Racon and the original reads by using neural network model to achieve high-quality polished assemblies (Oxford Nanopore Technologies Ltd. 2018) [56]. Contigs were Blast matched to the 16S rRNA sequences of the isolated bacterial strains and selected for quality assessment. QUAST (Quality Assessment Tool for Genome Assemblies) was used for checking assembly quality (Gurevich et al., 2013) [57]. The total length of ONT-sequenced genome was compared with a reference genome. Total length of ONT-sequenced genome with 90% of that of a reference genome was considered as a near-complete genome assembly, while lower than 90% was considered as a fragmented assembly. Prodigal was conducted to identify protein-coding genes and producing annotation files that describe gene locations and sequences for further analysis (Blast match with reference enzymes with hormone-metabolizing functions) (Hyatt et a., 2010) [58].

### Functional genomic analysis

In total, we isolated 55 gut bacterial strains with testosterone/estradiol metabolism activity from the eels. Among these, 22 strains were identified as known bacterial species (≥98.7% identity with a single species match per isolate). We identified *Deinococcu*s in the eel gut and purchased 11 *Deinococcus* species from BCRC. Bacterial genomes from Actinomycetes (n = 19), Bacilli (n = 2), Deinococci (n = 11), Alphaproteobacteria (n = 15), and Gammaproteobacteria (n = 13) were retrieved from the NCBI database for comparative functional genomic analysis. Genomes were analyzed by local BLAST against reference sequences; characterized enzyme protein sequences involve in testosterone or estradiol metabolism. Reference sequences included: 1) 9,10-seco pathway: *17β-hsd* (AZI34912.1) of *Novosphingobium tardaugens* NBRC16725 (Ibero et al., 2020) [59], *ORF17* (WP_003078230.1) of *Comamonas testosteroni* DSM50244 (Horinouchi et al., 2003) [60], *kshA* (AAL96829.1) and *kshB* (AAL96830.1) of *Rhodococcus erythropolis* (van der Geize et al., 2002) [61]; 2) 4,5-seco pathway: *oecA* (ARS27503.1) of *Sphingomonas sp.* KC8 (Tian et al., 2022; Zhang et al., 2024) [62, 63], *aedA* (WP_213934927.1) and *aedB* (WP_213934929.1) of *Rhodococcus sp.* strain B50 (Hsiao et al, 2021) [64], and *edcA* (AZI36851.2) and *edcB* (AZI36859.1) of *Novosphingobium tardaugens* NBRC16725 (Ibero et al., 2020) [59]; 3) Estrogen → Androgen pathway: *emtA* (WP_145842117.1), *emtB* (WP_145842116.1) and *emtC* (WP_145842115.1) of *Denitratisoma sp.* strain DHT3 (Wang et al., 2020) [65]; 4) 2,3-seco pathway: *atcA* (AMN46163.1) of *Steroidobacter denitrificans* DSMZ18526 (Yang et al., 2016) [66] and *SRAH* (WP_067170571.1) of *Steroidobacter denitrificans* Chol-1S (Jacoby et al., 2025) [67]. BLAST results were visualized in a heatmap showing protein sequence identity between bacterial genomes and reference sequences. Color gradient (blue-white-red) indicates sequence identity. Protein sequences with ≥90% coverage (followed Duan et al. 2024) [68] are shown in bold text.

### Hormone Quantification

Plasma testosterone was quantified using the Testosterone Parameter Assay Kit (R&D Systems, Inc., USA). Undiluted plasma samples were analyzed following the manufacturer’s protocol. Absorbance was measured at 450 nm with 570 nm wavelength correction. Concentrations were calculated using a 4-parameter logistic (4PL) regression model with standards ranging from 0 to 10 ng/mL.

Plasma estradiol was measured using the Estradiol Parameter Assay Kit (R&D Systems, Inc., USA). Samples were diluted 1.6-fold according to protocol specifications. Absorbance was measured at 450 nm with 570 nm wavelength correction. Final concentrations were calculated using a 4PL regression model (standard range: 0 to 3 ng/mL) and adjusted for dilution factor.

Plasma 11-KT concentrations were quantified using the 11-Ketotestosterone Competitive ELISA kit (Invitrogen, USA) with modifications. Briefly, 500 μl ethyl acetate (EA) was added to each 100 μl plasma sample in a 1.5 mL tube and vortexed for 10 minutes. Samples were centrifuged for 5 seconds, kept on ice, and the upper layer extracted. This extraction was repeated once to improve efficiency. Upper layer extracts from each sample were combined in a single 1.5 mL tube. EA was vacuum-dried and reconstituted in 110 μl assay buffer. Reagents and samples were added to wells, and absorbance was measured at 450 nm. 11-KT concentrations were calculated using a standard curve (0 to 2 ng/mL).

### Statistical Analysis

Data normality was assessed using the Shapiro-Wilk test. Data with normal distributions were analyzed using Welch’s t-test; non-normal distributions using Wilcoxon rank sum test. Significance was set at *p* < 0.05. Bacterial analyses included: 1) Differential abundance: lefser (Khleborodova et al., 2024) based on LEfSe (Segata et al., 2011); 2) Alpha diversity: two-sample t-test; 3) Beta diversity: PERMANOVA for bacterial community structure.

## Results

### Morphometric and physiological characterization of developmental stages in captive *Anguilla bicolor pacifica*

Wild glass eels were collected from an estuary in Hualien (eastern Taiwan) and identified as *A. bicolor pacifica* through maximum likelihood phylogenetic analysis of *COI* sequences (**Supplemental Figure S1**). The collected glass eels were cultivated in a freshwater fish farm for three years, with individuals sampled (n ≥ 6 per stage) at key developmental milestones: glass eel (GL), elver (EV), sex-undetermined eels (SU), and sex-determined eels (male [M] and female [F]). Morphometric analysis demonstrated progressive increases in body dimensions throughout development. Body length increased significantly across all developmental stages (all pairwise comparisons *p* < 0.01), progressing from 4.8 ± 0.12 cm in glass eels to 14.2 ± 0.86 cm in elvers, 33.4 ± 0.59 cm in sex-undetermined eels, 45.4 ± 1.14 cm in males, and 74.0 ± 1.01 cm in females (**Figure 2A**). Body weight exhibited a parallel developmental trajectory: GL (0.1 ± 0.01 g), EV (4.6 ± 0.84 g), SU (68.4 ± 5.07 g), M (248.7 ± 20.98 g), and F (1271.2 ± 26.61 g), with all groups significantly different from one another (*p* < 0.01) (**Figure 2B**). Sexual dimorphism was evident in the gonadosomatic index (GSI), with females displaying a substantially higher index (2.7 ± 0.24%) compared to males (0.4 ± 0.06%, *p* < 0.001) (**Figure 2C**).

**Figure 2.**
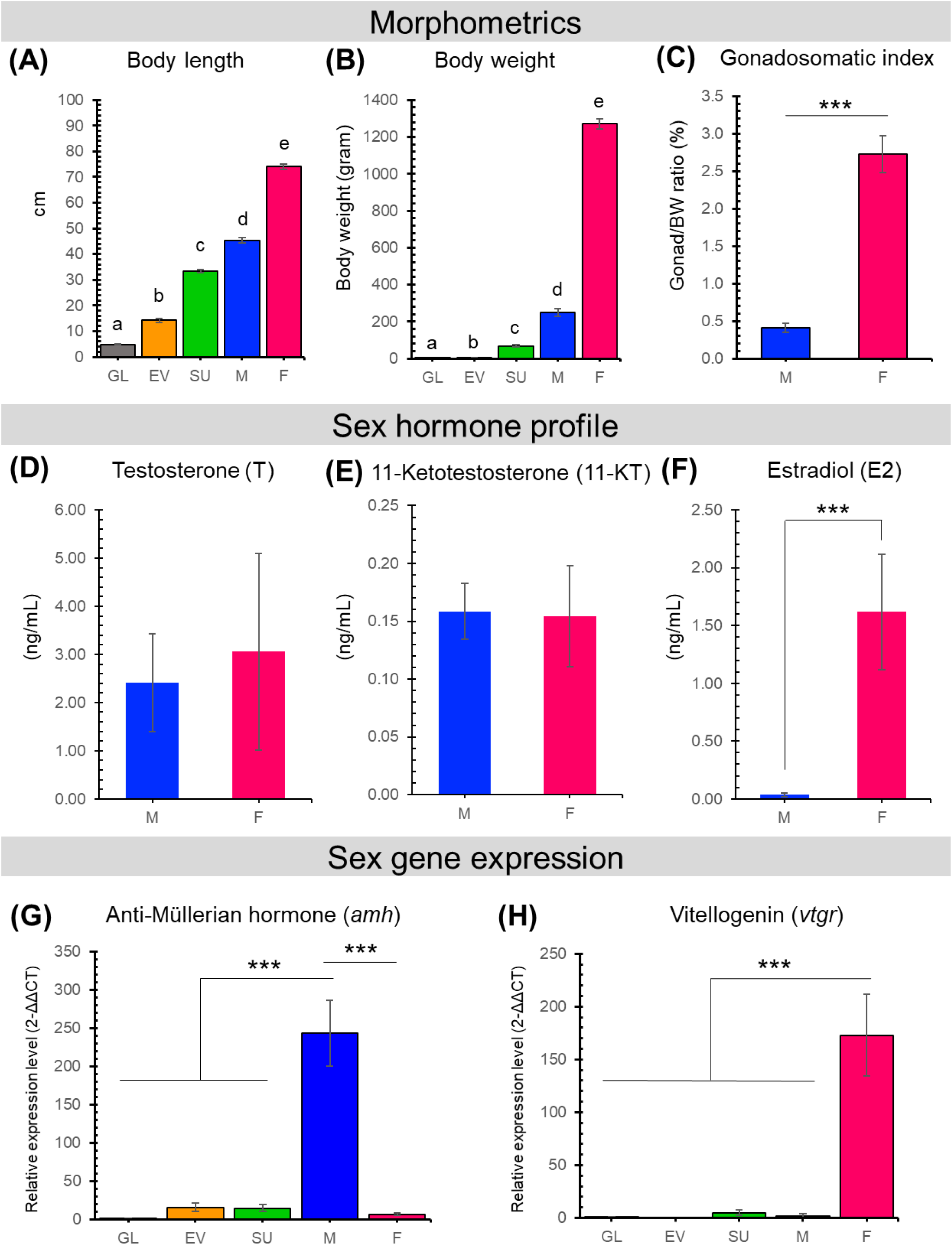
The morphometrics, sex hormone profiles, and expression of the selected sex-specific genes (*amh* and *vtgr* for male and female, respectively) of eel individuals collected at different development stages. The eel morphometrics determined in this present study include (**A**) body length (cm), (**B**) body weight (gram), and (**C**) Gonadosomatic index (GSI; %). Major sex hormones detected in the sex-determined eels include (**D**) testosterone, (**E**) 11-ketotestosterone, and (**F**) estradiol. Sex-specific genes studied in the eel samples include (**G**) *amh* and (**H**) *vtgr*. Number of eel individuals collected at different development stages: GL, n = 6; EV, n = 8; SU, n = 10; M, n = 10, and F, n = 10. Different letters (a∼e shown in Figures 2A and **2B**) indicate significant differences between groups (*p* < 0.01). Data with normal distributions were analyzed using Welch’s t-test; data with non-normal distributions were analyzed using the Wilcoxon rank-sum test; \*\*\**p* < 0.001.

Sex determination in adult eels was confirmed through integrated morphological assessment, genetic marker analysis, and hormonal profiling. Steroid hormone analysis revealed distinct profiles between sexes, with testosterone serving as the predominant sex steroid in both sex-determined males and females (mean serum concentrations > 2.0 ng/mL) (**Figure 2D**). Conversely, 11-ketotestosterone remained consistently low (< 0.2 ng/mL) across both sexes (**Figure 2E**). ELISA quantification revealed pronounced sex-specific differences in estradiol concentrations. Female eels exhibited markedly elevated estradiol levels (1.62 ± 0.50 ng/mL) compared to males (0.04 ± 0.02 ng/mL, *p* < 0.001) (**Figure 2F**). Despite this difference, testosterone concentrations remained comparable between sexes, with males showing 2.4 ± 1.02 ng/mL and females 3.1 ± 2.05 ng/mL (**Figure 2D**).

To further validate sex determination, reverse-transcription quantitative PCR analysis was performed targeting sex-specific molecular markers: anti-müllerian hormone (*amh*) as a male-specific marker (Lin et al., 2021) and vitellogenin receptor (*vtgr*) as a female-specific marker (Morini et al., 2020). Gene expression patterns provided clear discrimination between sexes and developmental stages. Male eels demonstrated markedly elevated *amh* expression (243.4 ± 43.16) compared to all other developmental groups: glass eels (1.0 ± 0.01), elvers (15.5 ± 0.84), sex-undetermined eels (14.7 ± 4.88), and females (6.7 ± 1.55, *p* < 0.001) (**Figure 2G**). In contrast, *vtgr* expression exhibited a distinct developmental progression, increasing from glass eels (1.0 ± 0.00) through elvers (0.1 ± 0.08) and sex-undetermined eels (5.2 ± 2.23), ultimately reaching peak expression in females (172.8 ± 38.69). Male eels maintained consistently low *vtgr* expression (2.2 ± 1.50, *p* < 0.001) (**Figure 2H**). This comprehensive analysis integrating morphometric data, gene expression profiles, and sex steroid concentrations establishes *amh* and *vtgr* as robust and reliable molecular markers for sex determination in male and female eels, respectively.

### Dynamic changes in gut microbial diversity of *Anguilla bicolor pacifica*

Gut microbiota composition was characterized across eel developmental stages using 16S rRNA gene amplicon sequencing (**Figure 3**). Following quality control, sequences were normalized to 2,499 reads per sample. Rarefaction analysis demonstrated that most eel samples reached saturation at approximately 600 reads, while feed pellet (FP) samples—the primary food source for elvers and adult eels—approached plateaus at the full 2,499 reads (**Supplemental Figure S2**). Shannon diversity index curves plateaued at 600 reads across all sample types, including environmental controls, confirming adequate sequencing depth.

**Figure 3.**
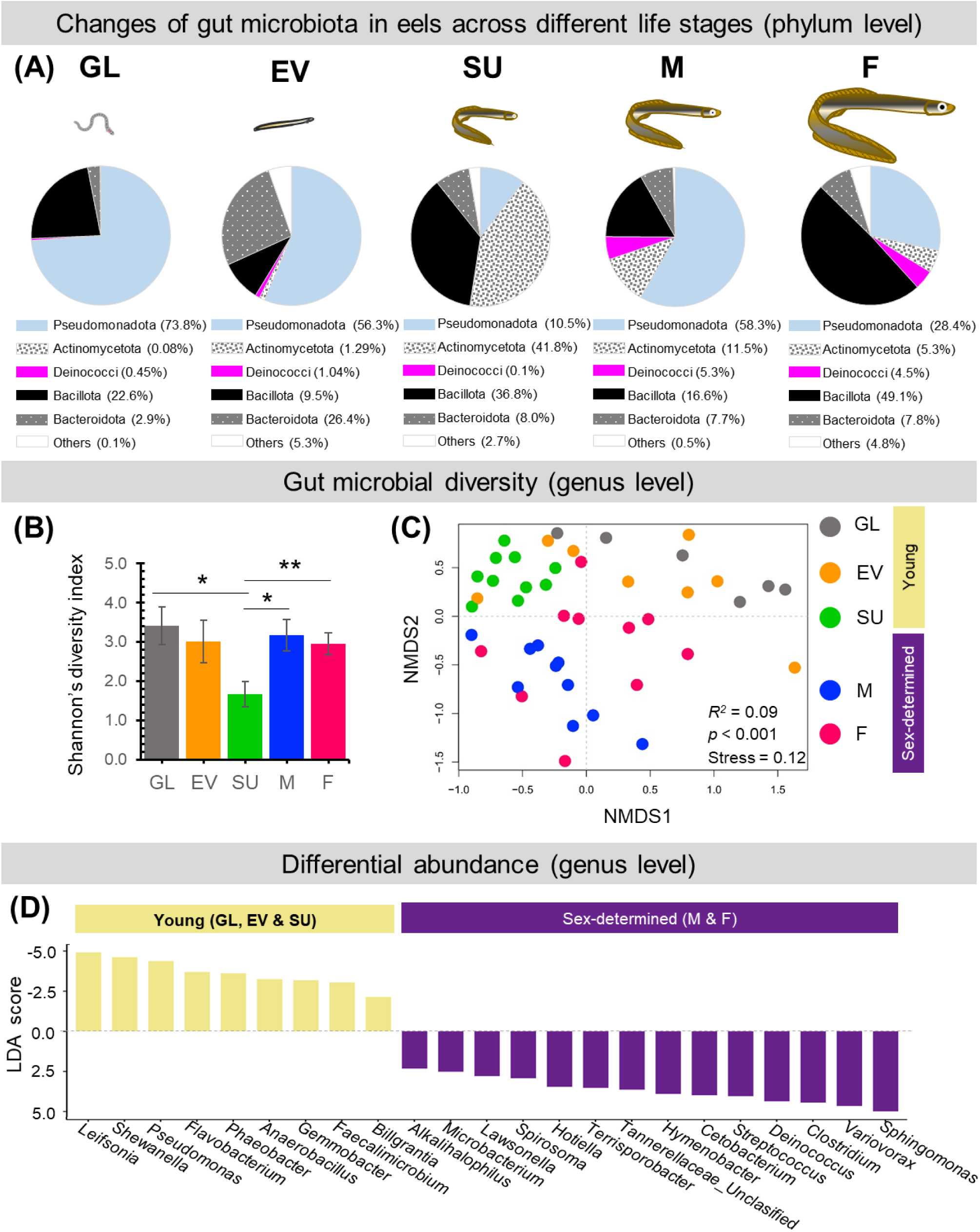
Changes of gut microbiota in eels across different life stages. (**A**) Pie charts show the dominant microbial phyla in the eel gut; (**B**) Shannon’s diversity index of the eel gut microbiota (genus level). Data with normal distributions were analyzed using Welch’s t-test; data with non-normal distributions were analyzed using the Wilcoxon rank-sum test; \**p* < 0.05; \*\**p* < 0.01; (**C**) Gut microbial composition analysis (genus level) based on Bray-Curtis dissimilarity and visualized in NMDS (PERMANOVA: *R^2^* = 0.09; *p* < 0.001); (**D**) Differential abundance of gut microbial genera (LEfSe). Barplot shows significantly dominant bacterial genera in either young (GL, EV, and SU; n = 24) or sex-determined adults (M and F; n = 20) groups. Genera are ranked by their LDA score.

Phylum-level analysis revealed dramatic microbial community restructuring throughout eel development (**Figure 3A**). Glass eel (GL) microbiomes were dominated by Pseudomonadota (73.8%), with Bacillota (22.6%) and Bacteroidota (2.9%) as minor constituents. During the elver (EV) stage, Pseudomonadota remained predominant (56.3%) but showed reduced abundance, coinciding with increased Bacteroidota representation (26.4%) and emergence of Spirochaetota (4.2%). Sex-undetermined eels (SU) exhibited the most pronounced taxonomic shift, characterized by Actinomycetota (41.8%) and Bacillota (36.8%) co-dominance, representing a departure from the Pseudomonadota-dominated profile of earlier stages. Adult stages showed sex-specific patterns: males (M) demonstrated renewed Pseudomonadota dominance (58.3%), while females (F) displayed peak Bacillota abundance (49.1%). Notably, Deinococcota showed progressive enrichment from juvenile to adult stages, contrasting with the early-stage dominance of Pseudomonadota (**Figure 3A**).

Alpha diversity, measured by Shannon’s index at the genus level, exhibited stage-specific fluctuations: GL (3.4 ± 0.484), EV (3.0 ± 0.538), SU (1.7 ± 0.327), M (3.168 ± 0.399), and F (2.9 ± 0.277) (**Figure 3B**). The marked reduction in diversity during the SU stage suggests a bottleneck effect during sexual maturation. Beta diversity analysis using non-metric multidimensional scaling (NMDS) based on Bray-Curtis dissimilarity revealed distinct community structures across developmental stages. NMDS ordination demonstrated a good fit (stress = 0.12), and PERMANOVA confirmed significant differences among life stages (*R²*= 0.09, *p* < 0.001) (Figure 3C). Integration of environmental controls demonstrated that eel gut communities formed discrete clusters, clearly separated from surrounding freshwater, red worms (primary food source for glass eels), and feed pellets (PERMANOVA: *R²*= 0.07, *p* < 0.001) (**Supplemental Figure S3**). Pairwise comparisons between consecutive developmental stages confirmed significant community transitions: GL-EV (*R²*= 0.23, *p* < 0.001), EV-SU (*R²*= 0.18, *P* < 0.001), SU-M (*R²*= 0.24, *p* < 0.001), SU-F (R²= 0.12, *p* = 0.006), and M-F (*R²*= 0.12, *p* = 0.017), indicating continuous microbiome restructuring throughout development (**Supplemental Figure S4**).

Linear discriminant analysis effect size (LEfSe) identified distinctive bacterial genera distinguishing sex-determined eels (M & F) from younger eels (GL, EV & SU) (**Figure 3D**). Sex-determined eels harbored 13 significantly enriched bacterial genera, with *Sphingomonas* showing the highest discriminatory power (LDA score: -4.99). Other notably enriched adult-associated genera included *Variovorax* (-4.66), *Clostridium* (-4.44), *Deinococcus* (-4.37), and *Streptococcus* (-4.04). Conversely, younger stages were characterized by nine significantly enriched genera (*p* < 0.05; LDA score > 2). *Leifsonia* exhibited the strongest juvenile association (LDA score: 4.92), followed by *Shewanella* (4.62), *Pseudomonas* (4.38), and *Flavobacterium* (3.70). Further analysis revealed sex-specific microbial signatures in sex-determined eels (**Supplemental Figure S5**). Males showed significant enrichment of six bacterial species (*p* < 0.05; LDA score > 2), including *Sphingomonas glacialis* (LDA score: 5.23), *Variovorax rhizosphaerae* (4.84), and *Deinococcus aquiradiocola* (4.28). Females exhibited enrichment of three species: *Staphylococcus warneri* (-4.20), *Deinococcus geothermalis* (-4.17), and *Anoxybacillus kestanbolensis* (-3.36).

Venn diagram analysis identified five core bacterial genera persistently present across all developmental stages: *Deinococcus*, *Pseudomonas*, *Acinetobacter*, *Clostridium*, and *Aeromonas* (**Figure 4A**). Among these core taxa, *Pseudomonas* demonstrated a progressive decline in relative abundance from glass eel to sex-determined eels (**Figure 4B1**), while *Deinococcus* exhibited the opposite trajectory, increasing throughout development (**Figure 4B2**). Bacterial richness (number of genera) was highest in adult females (112 unique genera), followed by adult males (89), elvers (67), glass eels (36), and sex-undetermined eels (32) (**Figure 4A**). This pattern suggests increasing microbiome complexity and specialization with sexual differentiation and adult development.

**Figure 4.**
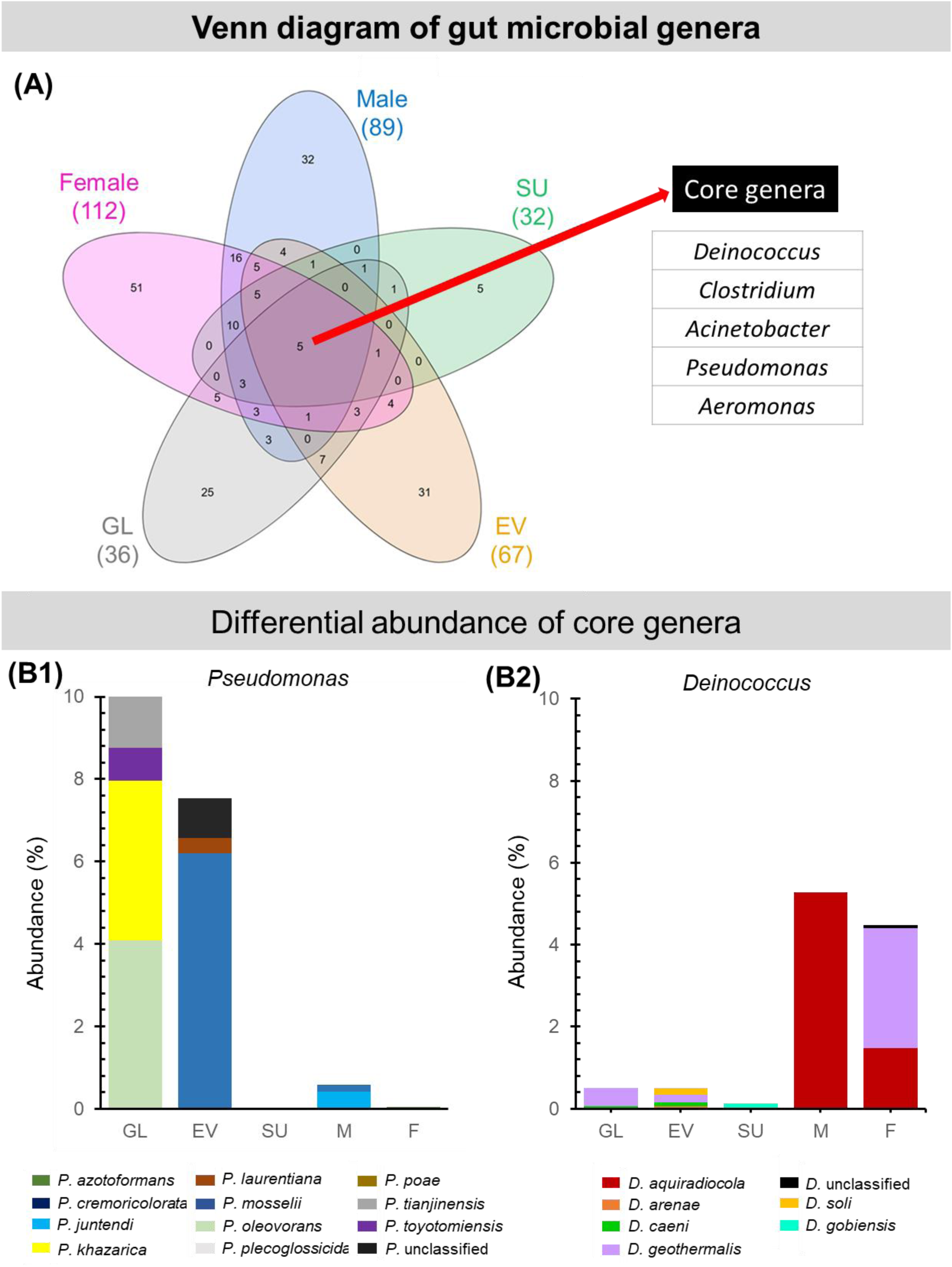
(**A**) Venn diagram analysis of gut microbiota identified five core genera (*Acinetobacter*, *Aeromonas*, *Clostridium*, *Deinococcus*, and *Pseudomonas*) shared across different eel life stages, with sex-determined adults showing higher bacterial richness (number of genera) than younger eels. (**B**) Relative abundance changes of two representative core genera across eel development. (**B1**) *Pseudomonas* demonstrated a progressive decline in relative abundance from glass eel to adult stages. (**B2**) *Deinococcus* showed an increasing abundance trend throughout eel development from glass eel to adult stages.

### Microbial interaction networks across eel life stages

The overall topology of microbial interaction networks at the genus level across eel life stages (GL, EV, SU, M, and F) is presented in Supplemental **Figure S6**. Centrality metrics identified the bacterial genus *Variovorax* as the most influential node (Rank 1: eigenvector = 1.000, degree = 5, closeness = 0.857, betweenness = 8.5), followed by *Deinococcus* (Rank 2: eigenvector = 0.782, degree = 3, closeness = 0.667, betweenness = 1.0), *Sphingomonas* (Rank 3) and *Rhodococcus* (Rank 4) (**Figures 5** and Supplemental **Table S1**). Detailed network analysis revealed two distinct major subnetworks. The first subnetwork, representing younger life stages (GL, EV, and SU), consisted primarily of five genera of the bacterial phylum Pseudomonadota (*Agrobacterium*, *Phaeobacter*, *Pseudomonas*, *Shewanella*, and *Vibrio*) and two genera of Bacillota (*Anaerostipes* and *Enterococcus*), with *Shewanella* as the key genus (**Figure 5**, left panel). The second subnetwork, representing mature life stages (F and M), was characterized by *Variovorax* as the key genus, forming significant connections with *Deinococcus*, *Hymenobacter*, *Methylobacterium*, *Nocardia*, *Rhodococcus*, and *Sphingomonas* (**Figure 5**, right panel). Correlation analysis revealed significant relationships between sex-determining genes and specific gut bacteria abundance at the genus level. The anti-müllerian hormone gene (*amh*) showed positive correlations with *Sphingomonas* (r = 0.40, *p* = 0.001***), *Variovorax* (r = 0.36, *p* = 0.002**), and a weak positive correlation with *Deinococcus* (r = 0.21, *p* = 0.046*). The vitellogenin receptor gene (*vtgr*) correlated positively with *Cetobacterium* (r = 0.211, *p* = 0.03*) and showed a marginally significant positive correlation with *Staphylococcus* (r = 0.21, *p* = 0.06) (Supplemental **Table S2**).

**Figure 5.**
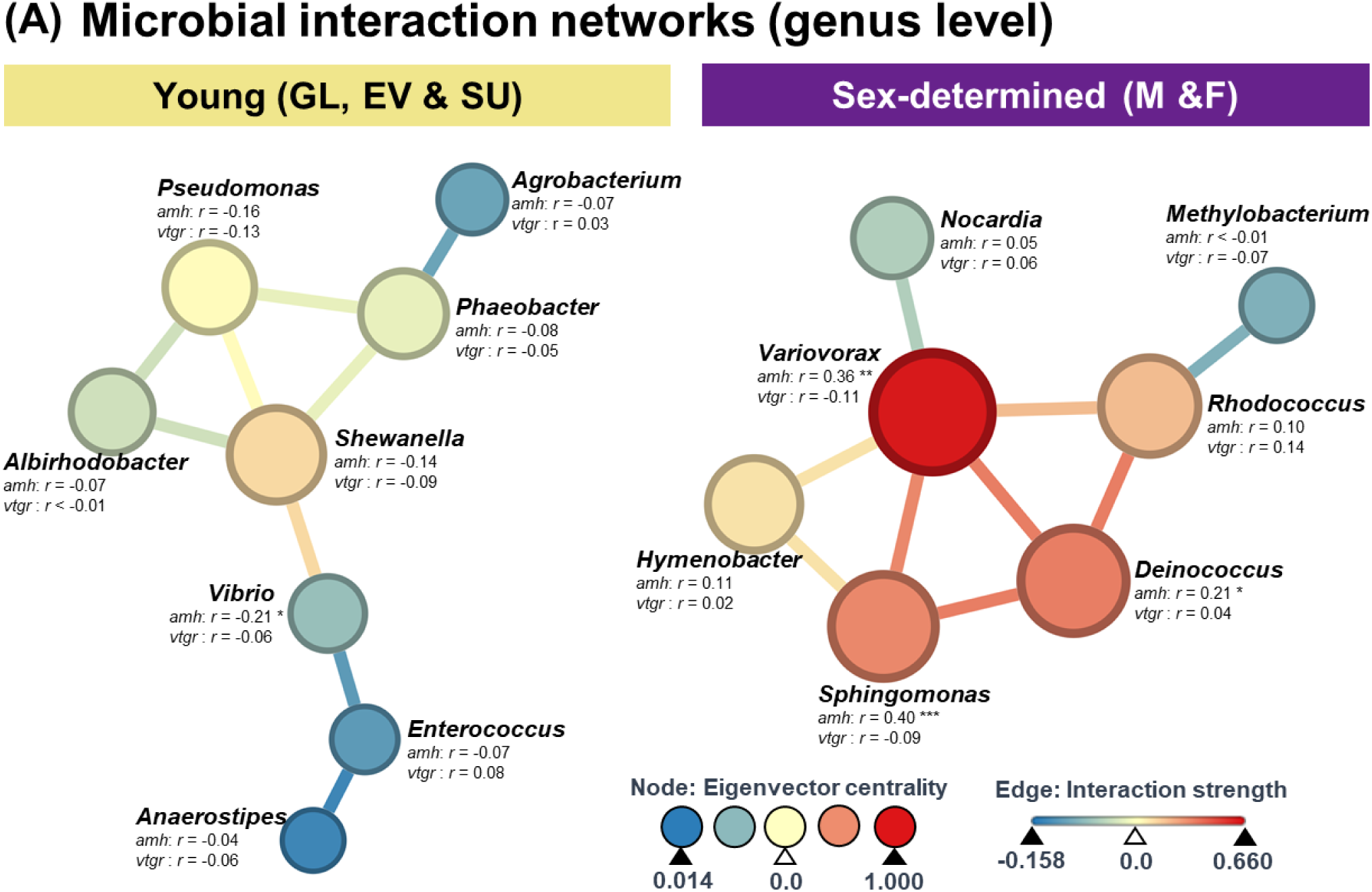
Microbial interaction subnetworks at the genus level across eel life stages and correlations between microbial genera and sex-specific gene expression. Each node represents a microbial genus, with node color and size indicating the eigenvector centrality score. Edge color represents the interaction strength between nodes, reflecting correlations between microbial genera. Correlations were calculated using FastSpar with 1000 bootstrap iterations. Correlations between microbial genera and the expression of *amh* and *vtgr* are indicated as correlation coefficients (r values). Asterisks indicate significance levels for correlations between microbial genera and the expression of *amh* and *vtgr*: \**p* < 0.05; \*\**p* < 0.01; \*\*\**p* < 0.001.

### Culture-dependent identification of sex steroid-metabolizing gut microbes

LEfSe and network analysis revealed several bacterial genera associated with mature eels, potentially linked to host hormone production. To validate their sex steroids-metabolizing function, we cultivated eel gut microbiota and employed culture-dependent approaches. We tested microbial steroid metabolism under strictly anaerobic, denitrifying, and microaerobic conditions. No sex steroids-metabolizing anaerobes were identified from the eel gut microbiota. By contrast, we collected 66 bacterial strains, all of which were capable of metabolizing sex steroids under microaerobic conditions (**Table 1** and Supplemental **Dataset S1**).

**Table 1.** Sex hormone-metabolizing bacteria (55 strains isolated from eel gut microbiome and 11 commercially purchased *Deinococcus* strains). Individual bacterial strains were identified by 16S rRNA gene sequencing and BLAST analysis against NCBI reference genomes. Steroid metabolism was evaluated using phosphate-buffered chemically defined medium containing 1 mM testosterone, 11-ketotestosterone, or estradiol as the sole carbon source. Strain-level data are provided in **Dataset S1**. Abbreviations: NA, no activity; NP, no product detected; UC, uncharacterized product; AD, 4-androstene-3,17-dione; ADD, androsta-1,4-diene-3,17-dione; DT, 1-dehydrotestosterone; E1, estrone.

| Number of bacterial isolates | Bacterial genera | Bacterial class | Major testosterone metabolites | Major 11-Ketotestosterone metabolites | Major estradiol metabolites |
| --- | --- | --- | --- | --- | --- |
| 7 | *Gordonia* | Actinomycetes | AD, ADD, NP, UC | NP, UC | E1, NP |
| 1 | *Microbacterium* | Actinomycetes | AD | UC | E1 |
| 2 | *Mycobacterium* | Actinomycetes | NP | NP | E1 |
| 5 | *Mycolicibacterium* | Actinomycetes | NP | NP, UC | E1, NP |
| 4 | *Nocardia* | Actinomycetes | NP | NP, UC | E1, NP |
| 2 | *Rhodococcus* | Actinomycetes | NP | NP | E1 |
| 1 | *Cytobacillus* | Bacilli | AD | NA | E1 |
| 1 | *Paenibacillus* | Bacilli | AD | UC | E1 |
| 11 | *Deinococcus* | Deinococci | AD, UC | NP | E1 |
| 2 | *Brucella* | Alphaproteobacteria | AD | NP | E1, UC |
| 1 | *Devosia* | Alphaproteobacteria | NP | NA | NP |
| 1 | *Neotabrizicola* | Alphaproteobacteria | AD | NA | NA |
| 2 | *Paracoccus* | Alphaproteobacteria | AD | UC | E1 |
| 2 | *Qipengyuania* | Alphaproteobacteria | NP | NP | NP |
| 5 | *Sphingomonas* | Alphaproteobacteria | NP | NP | E1, NP |
| 1 | *Stappia* | Alphaproteobacteria | NA | NP | E1 |
| 1 | *Ectopseudomonas* | Gammaproteobacteria | NA | NA | NP |
| 1 | *Metapseudomonas* | Gammaproteobacteria | NP | UC | E1 |
| 13 | *Pseudomonas* | Gammaproteobacteria | AD, ADD, DT, NP | UC | E1, NP, UC |
| 1 | *Pseudoxanthomonas* | Gammaproteobacteria | AD | UC | NP |
| 1 | *Shewanella* | Gammaproteobacteria | AD | UC | E1 |
| 1 | *Stutzerimonas* | Gammaproteobacteria | NA | NA | E1, UC |

These sex steroids-metabolizing bacterial isolates belonged to 22 bacterial genera: *Brucella, Cytobacillus, Deinococcus, Devosia, Ectopseudomonas, Gordonia, Metapseudomonas, Microbacterium, Mycobacterium, Mycolicibacterium, Neotabrizicola, Nocardia, Paenibacillus, Paracoccus, Pseudomonas, Pseudoxanthomonas, Qipengyuania, Rhodococcus, Shewanella, Sphingomonas, Stappia, Stutzerimonas.* To elucidate their steroid metabolic pathways, we cultivated the bacterial isolates with three major sex steroids (testosterone, 11-ketotestosterone and estradiol) under microaerobic conditions. After 3-day cultivation, we extracted microbial products from bacterial cultures using ethyl acetate and analyzed the extracts through UPLC-HRMS.

Steroid profile analysis revealed that most bacterial isolates performed oxidation reactions on the C-1/C-2 and/or 17-hydroxyl groups of the three sex steroids, maintaining the intact four-ring structure. The major microbial products included 4-androsten-3,17-dione (AD), androsta-1,4-diene-3,17-dione (ADD), 1-dihydrotestosterone (DT), and estrone (E1). A subset of gut microbes (*Brucella* sp. C40 *Deinococcus ficus* CC-FR2-10, *Gordonia* sp. OP42R, *Gordonia* sp. OP52R, *Metapseudomonas lalkuanensis* T46, *Microbacterium* sp. 4w1, *Mycolicibacterium fortuitum* subsp. *fortuitum* OP41W, *Nocardia asteroides* OT71, *Nocardia* sp. ST81, *Paenibacillus lautus* P8, *Paracoccus shanxieyensis* C32, *Pseudomonas farsensis* T49, *Pseudomonas guariconensis* P2, *Pseudomonas guariconensis* P3, *Pseudomonas nicosulfuronedens* T16, *Pseudomonas nitritireducens* T47, *Pseudomonas* sp. T19, *Pseudoxanthomonas mexicana* C34, *Shewanella xiamenensis* P7-3, and *Stutzerimonas frequens* OP54R) partially degraded testosterone, 11-ketotestosterone and/or estradiol through unknown pathways, resulting in the accumulation of uncharacterized products (UC). Certain species (marked in red in supplemental **Dataset S1**; examples including *Brucella* sp. OT17, *Deinococcus ficus* CC-FR2-10, *Devosia* sp. E27-1-2, *Ectopseudomonas mendocina* SP16, *Gordonia* spp., *Metapseudomonas lalkuanensis* T46, *Mycobacterium* spp., *Mycolicibacterium* spp., *Nocardia* spp., *Pseudomonas* spp., *Pseudoxanthomonas mexicana* C34, *Qipengyuania* spp., *Rhodococcus* spp., *Sphingomonas* spp., *Stappia indica* OT9) completely degraded the tested steroids without accumulating detectable products (NP; no accumulated products) in the ethyl acetate extracts. Notably, numerous bacterial species from the taxa Actinomycetes and Alphaproteobacteria demonstrate the capacity to completely degrade 11-ketotestosterone, the primary androgenic hormone in teleost fish. In contrast, most bacterial isolates belonging to Gammaproteobacteria can metabolize testosterone but lack the enzymatic capability to degrade 11-ketotestosterone or estradiol.

### Functional genomics reveals the prevalence of steroid catabolic genes among eel gut bacterial isolates

Steroid degradation capabilities in eel gut microbiota are primarily limited to Actinomycetes, Alphaproteobacteria, and Gammaproteobacteria. These bacterial taxa employ oxygen-dependent pathways for steroid degradation that have been characterized at the molecular level. Despite low sequence homology (deduced amino acid sequence identities < 40%), these bacteria utilize the oxygen-dependent 4,5-seco and 9,10-seco pathways to degrade estradiol and testosterone, respectively (**Figure 6A**).

**Figure 6.**
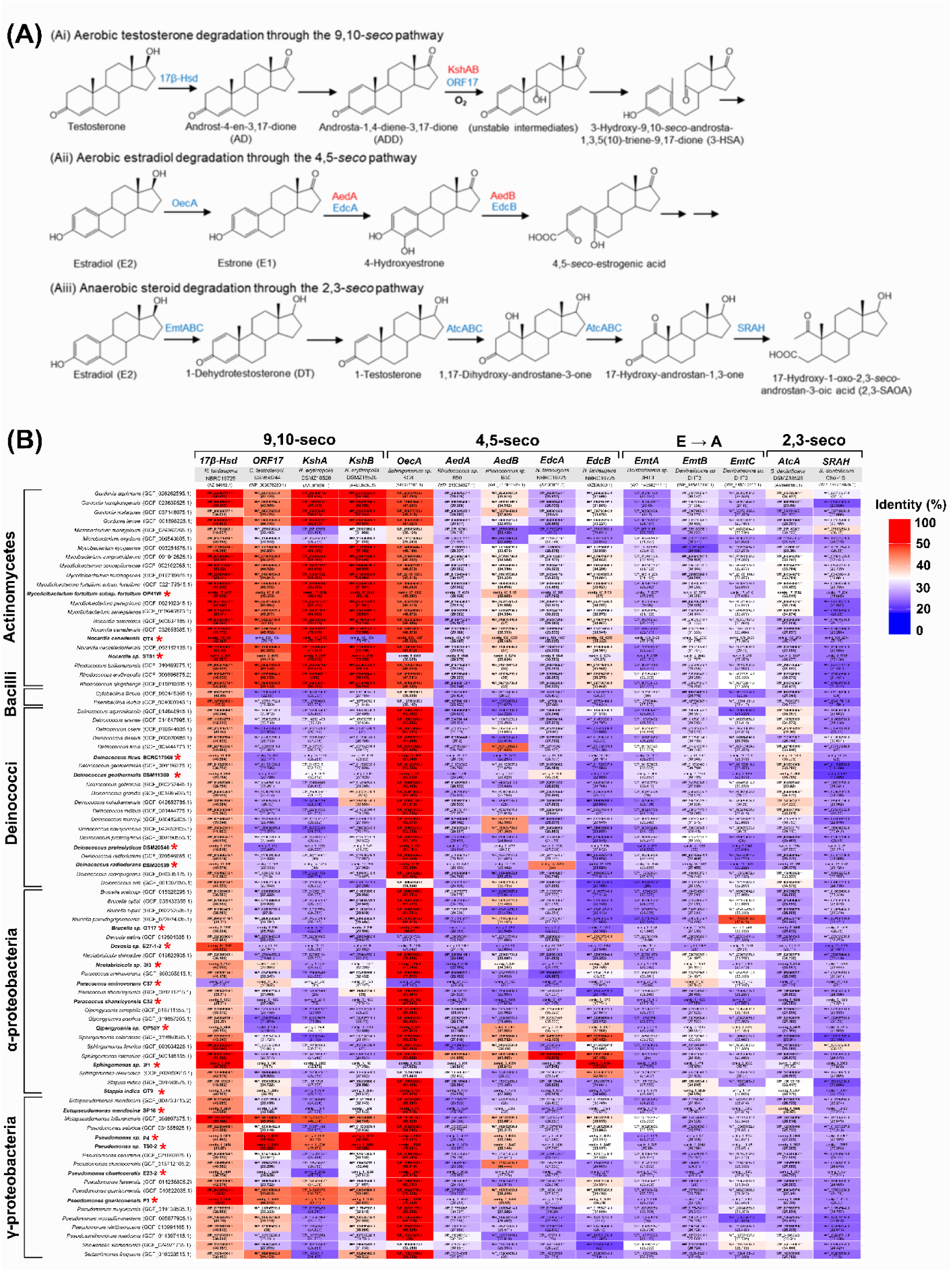
Distribution of steroid hormone metabolism genes in bacterial isolates from *Anguilla bicolor pacifica*. **(A) Established degradation pathways for testosterone and estradiol in bacteria.** (**Ai**) Aerobic testosterone degradation via the 9,10-*seco* pathway; (**Aii**) Aerobic estradiol degradation via the 4,5-*seco* pathway; (**Aiii**) Anaerobic steroid degradation via the 2,3-*seco* pathway. Red and blue annotations indicate enzymes from actinobacteria and proteobacteria, respectively. **(B) Gene prevalence heatmap across selected bacterial genomes.** Steroid metabolism genes were identified using BLAST searches with sequences from biochemically characterized enzymes or putative genes identified through comparative genomics. Reference sequences (deduced amino acid sequences): 9,10-*seco* pathway genes include 17*β*-*hsd* (AZI34912.1) from *Novosphingobium tardaugens* NBRC16725, ORF17 (WP_003078230.1) from *Comamonas testosteroni* DSM50244, and *kshA*/*kshB* (AAL96829.1/AAL96830.1) from *Rhodococcus erythropolis*; 4,5-*seco* pathway genes include *oecA* (ARS27503.1) from *Sphingomonas* sp. KC8, *aedA*/*aedB* (WP_213934927.1/WP_213934929.1) from *Rhodococcus* sp. B50, and *edcA*/*edcB* (AZI36851.2/AZI36859.1) from *N. tardaugens* NBRC16725; anaerobic estrogen-to-androgen biotransformation genes include *emtA*/*B*/*C* (WP_145842117.1/WP_145842116.1/WP_145842115.1) from *Denitratisoma* sp. DHT3; and 2,3-seco pathway genes include *atcA* (AMN46163.1) from *Steroidobacter denitrificans* DSMZ18526 and *SRAH* (WP_067170571.1) from *S. denitrificans* Chol-1S. The color gradient (blue-white-red) represents sequence identity percentage. Bold text indicates protein sequences with ≥90% coverage. Red asterisks indicate genomes of eel gut isolates sequenced in this study, while unmarked genomes represent closely related species obtained from NCBI with high 16S rRNA gene identity (96-100%) to the identified eel gut species.

We assessed the distribution of characterized steroid metabolism genes across genomes from isolated eel gut microbes (highlighted in bold) and closely related species (16S rRNA sequence identity >99%) (**Figure 6B**). The *17β-hsd* gene, encoding 17*β*-hydroxysteroid dehydrogenase, shows widespread distribution across these genomes. Actinobacteria and Proteobacteria utilize this enzyme to convert testosterone to androstenedione (AD) (Holert et al., 2018; Chiang et al., 2020) [69, 70]. We identified homologous genes in Deinococci species, whose gene products demonstrate highest sequence similarity to proteobacterial enzymes (up to 59.1%) and actinomycetes enzymes (up to 52.6%), with moderate similarity to Bacilli enzymes (up to 47.2%). In contrast, genes involved in androgenic core-ring degradation (including *ORF17*, *kshA*, and *kshB*) show limited distribution, primarily confined to Proteobacteria and Actinomycetes. Among Deinococci, these genes were identified only in select species such as *Deinococcus grandis* and *D. hohokamensis*.

For estradiol degradation, both Actinobacteria (Hsiao et al., 2021, 2022) [50, 64] and Alphaproteobacteria (Chen et al., 2017; Wu et al., 2019) [49, 71] employ the 4,5-*seco* pathway. The *oecA* gene, involved in converting estradiol to estrone, shows broad distribution across bacterial genomes, including Deinococci (with deduced amino acid sequence identity up to 53.4%). However, genes encoding estrogenic A-ring degradation enzymes (*aedA*, *aedB*, *edcA*, and *edcB*) are restricted to Alphaproteobacterial and Actinobacterial genomes (**Figure 6B**), suggesting that Bacilli and Deinococci species lack aerobic estrogen degradation capabilities.

Anaerobic degradation of sex steroids has been documented in only a few Beta- and Gamma-proteobacteria (Chiang et al., 2020; Wang et al., 2020) [65, 70]. Using characterized degradation genes from *Denitratisoma* sp. strain DHT3 as queries, we detected limited distribution of these genes among eel gut bacteria. Our comparative genomic analysis thus indicates that anaerobic steroid metabolism is not prevalent in the eel gut microbiome.

## Discussion

In this present study, we specifically investigated gut microbiota changes during sexual differentiation rather than sexual maturation (silvering) for three key reasons. First, silvering involves dramatic gut regression and atrophy as eels prepare for non-feeding oceanic migration, fundamentally altering the intestinal environment (Sudo et al., 2012) [19]. Second, the natural fasting behavior during this phase causes significant reductions in gut microbial diversity and density due to the absence of dietary substrates and deteriorating intestinal habitat. Third, the functional significance of gut microbiota in host physiology becomes negligible during silvering, as eels rely entirely on stored energy rather than active digestion and metabolism. By examining actively feeding, sexually differentiating eels, we captured biologically relevant host-microbiota interactions at a critical life stage where these relationships are most functionally significant and experimentally informative.

### Sex steroid profiles in short-fin eels

Our study provides important insights into the sex steroid profiles of short-fin eels (*A. bicolor*) cultivated under controlled freshwater conditions. The results demonstrate that serum testosterone levels in male short-fin eels were comparable to those in mature females, while serum estradiol levels in females were significantly higher than in males. However, we observed notably limited production of 11-ketotestosterone in both mature male and female eels under these freshwater conditions. The low 11-ketotestosterone levels observed in our study likely reflect the artificial freshwater cultivation environment rather than a fundamental species difference from Japanese eels. 11-Ketotestosterone is well-established as the primary androgen in teleost fish, including multiple eel species (Borg, 1994; Lokman et al., 2007) [72, 73], and its production is typically enhanced during sexual maturation and environmental transitions (Palstra et al., 2005; Thomson-Laing et al., 2019) [74, 75]. In natural conditions, eels undergo complex hormonal changes during their catadromous migration from freshwater to marine environments for reproduction (Sudo & Yada et al., 2022; van Ginneken & Maes, 2005) [20, 76]. The artificially maintained freshwater conditions in our study may have suppressed the normal elevation of 11-ketotestosterone that would occur during natural sexual maturation and seawater adaptation.

Previous studies have demonstrated that environmental factors, particularly salinity changes, significantly influence steroid hormone production in eels (Quérat et al., 1987; Sudo et al., 2011; Palstra et al., 2005) [74, 77, 78]. The transition to seawater conditions typically stimulates gonadotropin release and subsequent steroidogenesis, including 11-ketotestosterone production in males (Sudo et al., 2011) [78]. Therefore, the steroid profiles observed in our freshwater-maintained eels likely represent an incomplete maturation state rather than the full reproductive hormone profile characteristic of naturally maturing eels. Comparative studies examining hormone profiles across different salinity conditions and maturation stages would be valuable for understanding the environmental regulation of eel reproductive endocrinology (van Ginneken & Maes, 2005; Sudo et al., 2011) [76, 78].

### Enterohepatic circulation of sex steroids in eels

Sex steroid levels and gonadosomatic index (GSI) vary significantly across vertebrate species. Our findings demonstrate that *A. bicolor* females exhibit substantially elevated GSI and estradiol levels compared to other vertebrates. Specifically, these eels show GSI values of 2.5–3.0% and E2 concentrations of 1.1–2.1 ng/mL, which are markedly higher than those observed in healthy female rats (GSI: 0.038%, E2: 0.012 ng/mL; Hazarika et al., 2023; Yuliawati et al., 2020) [79, 80] and female mice (GSI: ∼ 0.15%, E2: ∼0.01 ng/mL; Zheng et al., 2024; Constantin et al., 2025) [81, 82].

The anatomical differences between eels and mammals may contribute to these elevated hormone levels. While mammals possess complex digestive systems with distinct small and large intestine divisions, eels have relatively simple digestive tracts—essentially a tube extending from the esophagus to the anus. The gut microbial communities of *A. bicolor* further distinguish these eels from mammals. While human gut microbiota are dominated by anaerobic bacteria (primarily Bacteroidota and Bacillota), eel gut communities are predominantly composed of aerobic or microaerophilic organisms, reflecting the dominance of Pseudomonadota and Actinomycetota. Although some obligate and facultative anaerobes (Bacteroidota and Bacillota) are present in eels, they represent a smaller fraction of the total community. This aerobic bias is further supported by our inability to isolate any anaerobic sex hormone-metabolizing bacteria from eel gut samples. The eel intestine harbors a greater abundance of aerobic hormone-metabolizing bacteria, including *Deinococcus*, *Rhodococcus*, and *Sphingomonas*, compared to human and mouse gut microbial communities (Hsiao et al., 2023; Chen et al., 2024; Brandon-Mong et al., 2024bioRxiv) [23, 24, 83]. These specialized microbial communities actively metabolize the sex hormones produced by the eel’s enlarged gonads (testosterone, 11-ketotestosterone, and estradiol), generating various hormone metabolites such as androstenedione, androstenediol, estrone, and uncharacterized hormone metabolites. The unique anatomical features of the eel digestive system may facilitate enterohepatic circulation of these steroid compounds. The relatively simple gut architecture potentially allows bacterial hormone metabolites to cross the intestinal barrier more readily and enter systemic circulation. This hypothesis is supported by previous research demonstrating that steroids can traverse the fish intestinal barrier and participate in enterohepatic circulation (Castejón et al., 2021; Pelissero & Sumpter, 1992) [84, 85].

The elevated hormone levels, combined with the presence of specialized metabolizing bacteria and permeable gut architecture, suggest that eels may have evolved an efficient enterohepatic recycling system for sex steroids. Future investigations should focus on characterizing the biological activity of these bacterial hormone metabolites and elucidating their specific physiological roles in eel reproduction and development. Understanding this enterohepatic circulation pathway could provide valuable insights into the unique reproductive strategies of anguillid eels and their remarkable life cycle adaptations.

### Developmental changes in eel gut microbiota composition

Our multi-omics analyses reveal significant shifts in gut microbiota composition throughout eel development, with distinct patterns emerging across life stages and between sexes. Early developmental stages, including glass eels and elvers, are dominated by the bacterial phylum Pseudomonadota. In contrast, sex-determined male and female eels exhibit increased abundance of Deinococcota, indicating a major compositional transition during sexual development.

Notable sex-specific differences emerge in sex-determined eels. Female eels harbor higher proportions of facultative anaerobes, primarily from the phylum Bacillota, while males maintain elevated levels of aerobic and microaerophilic bacteria, predominantly Pseudomonadota. Key bacterial genera in sex-determined eels include *Variovorax*, *Deinococcus*, *Sphingomonas*, and *Rhodococcus*. Particularly intriguing is the positive correlation between the anti-Müllerian hormone gene (*amh*), a male-specific marker, and the abundance of *Sphingomonas*, *Variovorax*, and *Deinococcus*. This correlation suggests potential functional interactions between these bacterial taxa and eel sexual development pathways.

The observed microbiota shifts likely reflect changes in gut physiology during eel development. As eels grow, their intestinal length increases substantially (Kužir et al., 2012; Knutsen et al., 2021) [86, 87], potentially altering the gut microenvironment from subnormoxic to hypoxic conditions. This environmental transition subsequently influences both microbiota composition and metabolic functions. Our data demonstrate that glass eels, elvers, sex-undetermined eels, and male adults maintain guts dominated by aerobic and microaerophilic microbes. In contrast, female guts shift toward facultative anaerobe dominance, possibly reflecting the longer gut length and altered oxygen availability in reproductively active females. These findings align with previous research showing that Bacillota abundance positively correlates with fish body size, while Pseudomonadota abundance decreases in larger fish (Jiang et al., 2019) [88]. The progressive changes in gut oxygen availability during eel development appear to be key drivers of both microbial community structure and functional capacity, ultimately shaping the unique hormone-metabolizing capabilities observed in different life stages and sexes.

### Sex steroid metabolism by eel gut microbiota

Our study provides the first comprehensive characterization of sex steroid-metabolizing gut bacteria in teleost fish. We successfully isolated and identified 66 bacterial strains capable of metabolizing sex hormones, thereby validating the functional predictions derived from our multi-omics analyses. Multiple species from the genera *Rhodococcus* and *Sphingomonas* demonstrated robust aerobic steroid metabolism capabilities, which aligns with the high prevalence of steroid degradation genes detected in their genomes.

The eel gut microbiome’s steroid metabolism differs fundamentally from mammalian systems. Mammalian colons represent oxygen-depleted, highly reduced environments (approximately -200 mV) dominated by strictly anaerobic bacteria such as *Clostridium* species, which utilize sex steroids as electron acceptors (Chen et al., 2024) [24]. In contrast, the short intestinal tracts of eels create a microaerobic ecosystem that supports the activity of oxygenases and corresponding aerobic microorganisms. This unique environment enables complete aerobic degradation of estradiol, a process that is predominantly reported under oxygen-rich conditions.

Isolating bacterial strains capable of complete estradiol degradation as single cultures remains a significant challenge in environmental microbiology, particularly from complex matrices such as sewage, wastewater, and estuarine sediments. In sewage and wastewater systems, bacterial communities are typically dominated by Betaproteobacteria, followed by Gammaproteobacteria and Actinomycetes (Do et al., 2019) [89]. Estuarine environments show a different pattern, with Gammaproteobacteria predominating, followed by Alphaproteobacteria and Actinomycetes (Ghosh et al., 2019) [90]. Most bacteria isolated from these environments achieve only partial estradiol degradation, typically converting it to intermediate metabolites like estrone, which may persist and retain estrogenic activity (Chen et al., 2018; Yu et al., 2007) [91, 92]. For instance, activated sludge isolates frequently convert estradiol to estrone but fail to mineralize estrone further, with only rare strains demonstrating complete degradation and growth on estradiol as the sole carbon source (Yu et al., 2007) [92]. Furthermore, degradation efficiencies are often enhanced in bacterial consortia rather than single isolates, indicating that complete estradiol mineralization typically requires cooperative processes involving multiple microbial species with complementary metabolic capabilities (Chen et al., 2018; Li et al., 2018; Yu et al., 2007) [91–93].

The isolation of multiple bacterial genera from the eel gut capable of completely degrading estradiol as single isolates represents a significant finding that provides novel insights into microbial estradiol catabolism. The bacterial genera we isolated—*Gordonia*, *Mycolicibacterium*, *Nocardia* (Actinomycetes), *Devosia*, *Qipengyuania*, *Sphingomonas* (Alphaproteobacteria), and *Ectopseudomonas*, *Pseudomonas*, *Pseudoxanthomonas* (Gammaproteobacteria)—are consistent with taxa previously implicated in steroid degradation. However, these genera are rarely reported as single isolates capable of fully mineralizing estradiol (Chen et al., 2018; Ibero et al., 2020; Yu et al., 2007) [59, 91, 92]. The ability of these eel gut bacteria to utilize estradiol as a sole carbon and energy source suggests specialized metabolic adaptations. These adaptations may be linked to the unique ecological niche of the eel gut, where steroid hormones are likely present at elevated concentrations due to the host’s distinctive physiology and reproductive biology. This finding highlights the potential of fish gut microbiomes as sources of novel steroid-degrading bacteria with biotechnological applications.

### The gut microbe *Deinococcus* spp. as newly identified steroid metabolizers

A particularly significant finding in our study involves the physiological and molecular characterization of *Deinococcus* species from the eel gut, which showed increasing abundance from glass eel to adult stages. While *Deinococcus* spp. have been previously found in European eel guts and characterized as microaerophiles growing optimally in hypoxic environments (Huang et al., 2018; Liedert et al., 2012) [29, 94], our study provides the first evidence of their capability for aerobic steroid metabolism. *Deinococcus* belongs to phylum Deinococcota, comprising three genera: *Deinococcus*, *Deinobacterium*, and *Truepera* (Lum et al., 2024) [95]. As radiation-resistant extremophiles, these species have been isolated from diverse environments, including wastewater systems (Liu et al., 2023) [96]. Despite their ecological significance, steroid hormone metabolic pathways in *Deinococcus* remain poorly understood, with only *D. actinosclerus* previously identified as capable of partial estradiol oxidation (with estrone as the major product) via *oecA*-encoded 3*β*,17*β*-hydroxysteroid dehydrogenase (Xiong et al., 2018) [97]. Consistently, most *Deinococcus* species in our study simply oxidized the 17-hydroxy group of sex steroids, causing mild loss in hormonal activity. However, at least one species, namely *Deinococcus ficus*, exhibited remarkable differential metabolic capabilities toward structurally related androgens. While testosterone was only partially metabolized, producing androstenedione and uncharacterized steroidal metabolites, 11-ketotestosterone underwent complete degradation. This substrate specificity suggests distinct metabolic pathways for these androgens. This differential capability highlights the substrate-specific nature of bacterial steroid metabolism and suggests that 11-ketotestosterone may represent a preferred carbon source for this isolate, providing the first evidence of host-microbe interactions between eel sex hormones and steroid-metabolizing *Deinococcus* while advancing our understanding of sex hormone metabolism in eel-associated gut microbiota.

### Implications for host-microbe interactions and eel development

Our findings suggest that interactions between the eel gut microbiota and their hosts are bidirectional. The aerobic bacteria we identified (e.g., *Sphingomonas* and *Rhodococcus*) likely consume gut sex steroids, potentially decreasing circulating levels of androgens and estrogens in their hosts. This may negatively impact eel development and sexual maturation. As these bacterial species often inhabit aquatic environments and represent dominant microorganisms in aquaculture, removing sex steroid-degrading bacteria from the eel gut and surrounding environments may facilitate the development and sexual maturation of captive eels.

Recent studies using mouse models have demonstrated that testosterone-degrading bacteria can significantly decrease serum testosterone levels through enterohepatic circulation (Hsiao et al., 2023) [23]. Similarly, these accumulated bacteria might continually influence cultivated eels in aquaculture ponds, creating a feedback loop that affects hormonal balance and reproductive development. Conversely, wild eels might be less influenced by the same bacteria due to multiple environmental factors, such as salinity, temperature, and environmental hormones. For example, environmental hormones coexist in habitats of benthic eels. Higher concentrations of hormones may alter microbiota compositions and the population density of various hormone-degrading bacteria. Both environmental hormones and bacteria may influence benthic fishes’ growth and maturation in downstream rivers. This may explain why large variations in gut microbiota appear in aquatic animals, such as wild benthic eels.

## Conclusion

This study demonstrates that host-gut microbiota interactions play crucial roles in developmental programming and sexual differentiation of *A. bicolor*. The identification of novel steroid-metabolizing bacteria, particularly *Deinococcus* species with unique substrate specificities, expands our understanding of microbial steroid metabolism and host-microbe interactions in aquatic vertebrates. Through our multi-omics approach, we predicted the steroid hormone metabolizing function of several bacterial genera through correlation analyses and validated their functions through culture-dependent approaches. Microbial interaction network analysis shows that the abundance of *Deinococcus*, *Sphingomonas*, and *Variovorax* was positively correlated with each other and may interact for synergistic hormone metabolism. This metabolic specificity suggests the potential application of these bacterial genera as biomarkers for eel sexual development.

The isolation of multiple bacterial genera capable of completely degrading estradiol as single isolates has important biotechnological implications. These bacteria could be utilized for bioremediation of endocrine-disrupting compounds in aquatic environments or for developing novel bioprocesses for steroid hormone degradation. Additionally, understanding the specific mechanisms of steroid metabolism in these bacteria could inform strategies for managing hormone levels in aquaculture settings.

Further research is needed to: (i) establish the complete testosterone degradation pathway of *D. ficus* and identify corresponding degradation genes and enzymes; (ii) investigate the molecular-level interactions between sex steroid-metabolizing bacteria and circulating hormone profiles; (iii) validate whether transferring sex steroid-metabolizing bacteria, especially *D. ficus*, into the gut of elvers could delay sexual development; and (iv) conduct comparative studies of sex steroid profiles in different eel species under standardized environmental conditions to determine species-specific versus environmentally-induced hormonal differences. Additionally, future studies should examine the potential for manipulating gut microbiota composition to optimize eel reproduction in aquaculture settings. Understanding the precise mechanisms by which gut bacteria influence host hormone levels could lead to targeted interventions that enhance reproductive success in captive eel populations, contributing to conservation efforts for this endangered group of fish.

