## Supplemental figures and tables for "Multi-omics integration uncovers host-microbiota crosstalk underlying sexual differentiation in the shortfin eel *Anguilla bicolor pacifica*"

**This PDF file includes the following:**

Figs. S1 to S6
Tables S1 to S2

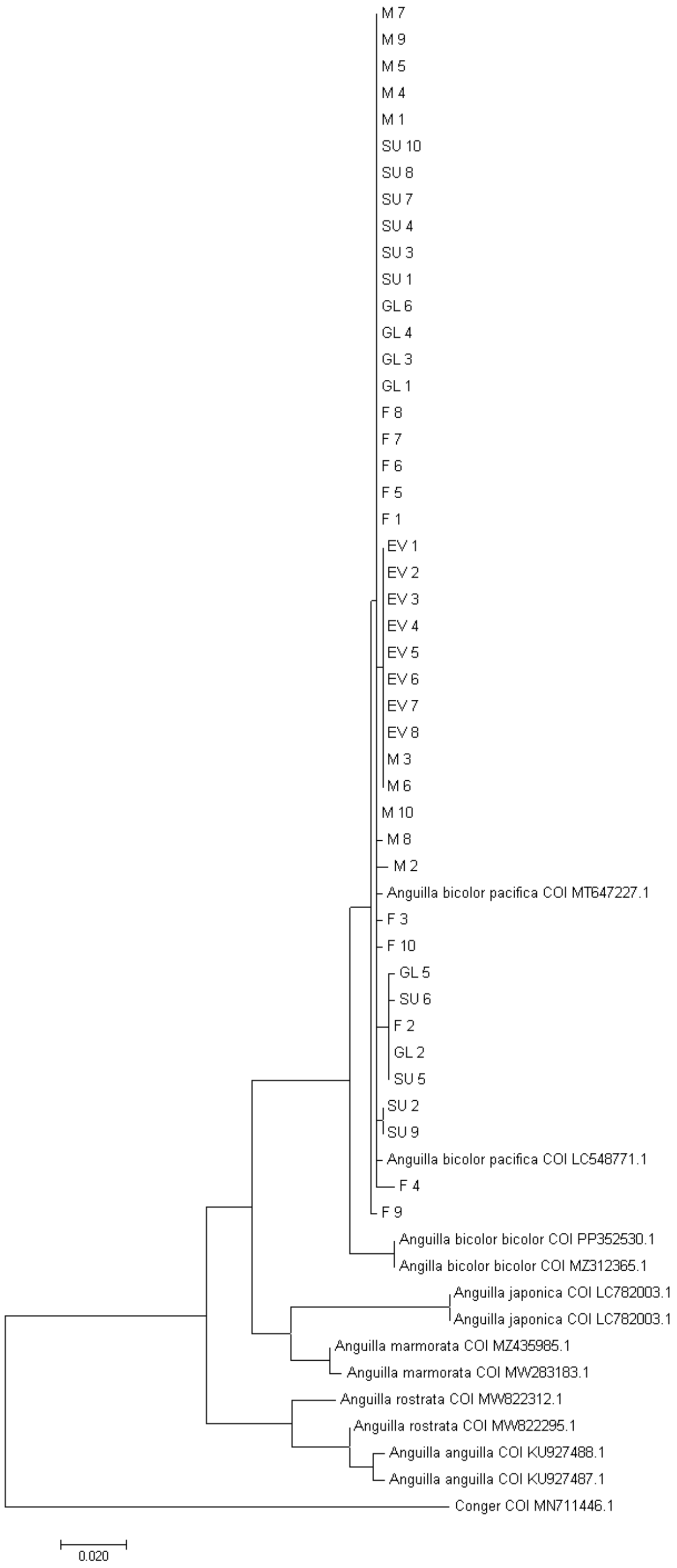


**Supplemental Figure 1.** Maximum likelihood phylogenetic tree based on *COI* sequences of *Anguilla bicolor pacifica* in this study and reference sequences. *Conger* served as an outgroup.

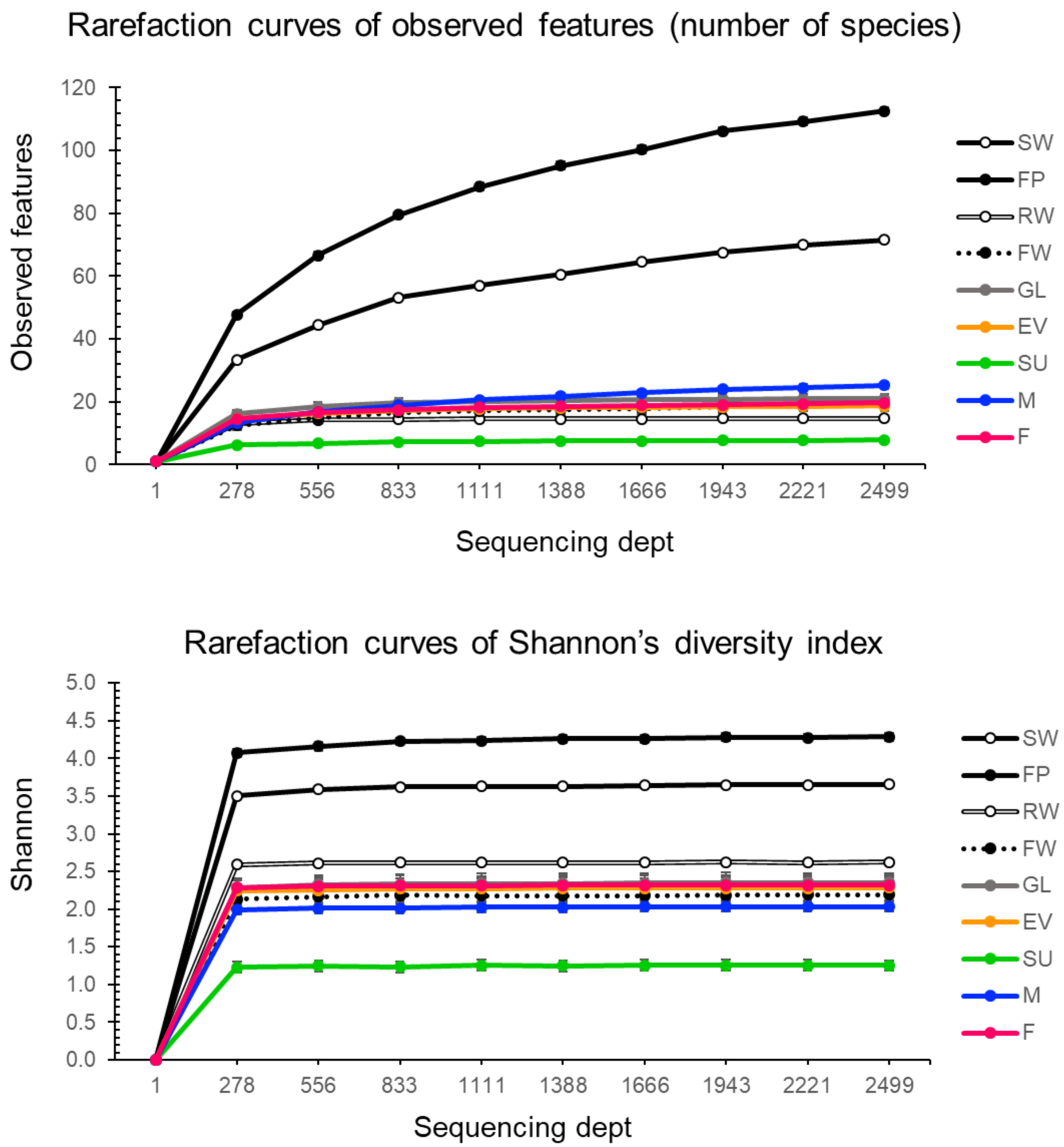


**Supplemental Figure 2.** Rarefaction curves of observed features (number of species) and Shannon's diversity index.

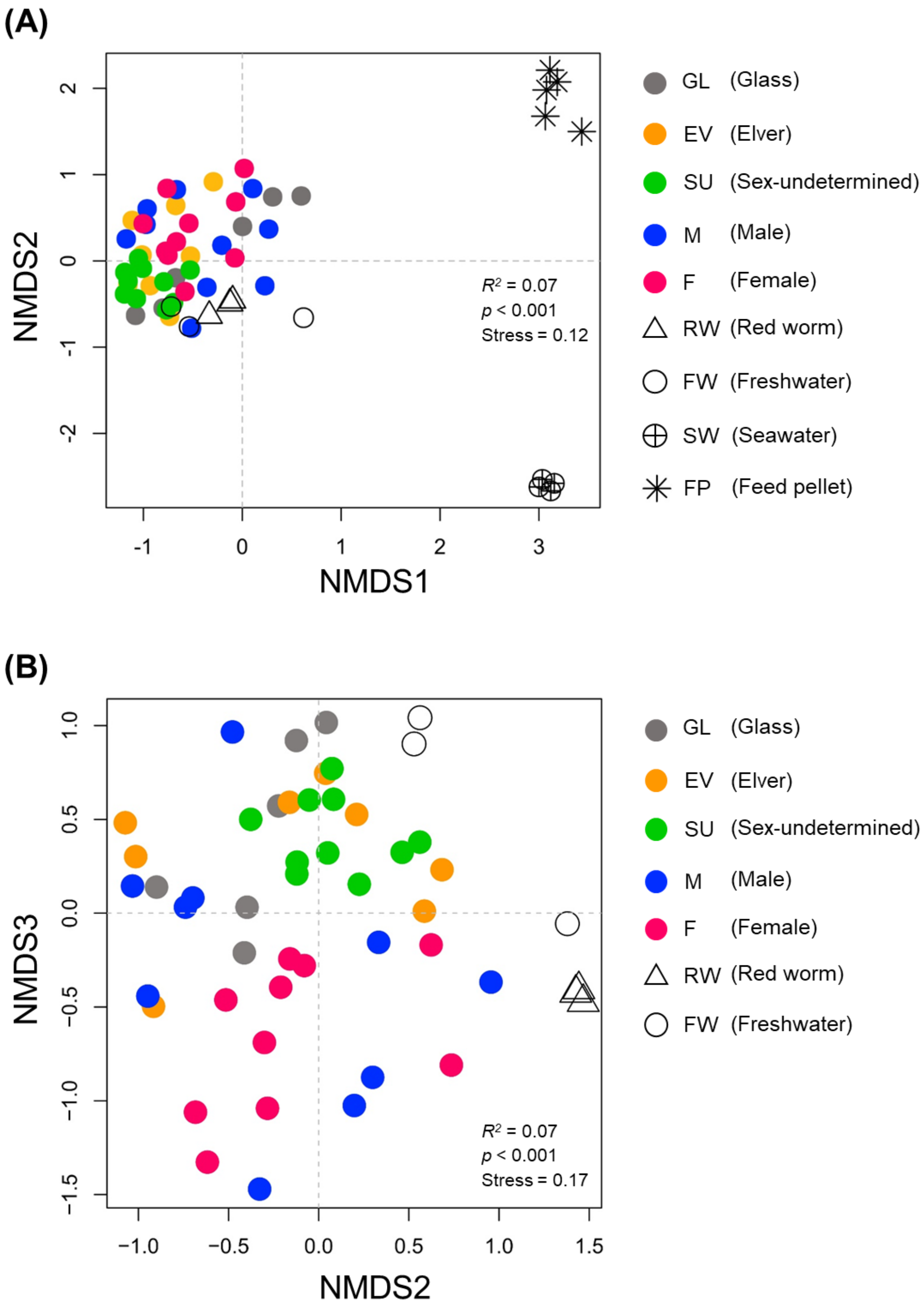


**Supplemental Figure 3.** Beta-diversity analysis of gut microbial communities and environmental controls at the genus level based on Bray-Curtis dissimilarity, visualized using NMDS. A) Gut microbial composition (GL, EV, SU, M, and F) and controls (RW: red worms; FW: freshwater; SW: seawater; FP: feed pellet). B) Gut microbial composition (GL, EV, SU, M, and F) with RW (red worms) and FW (freshwater).

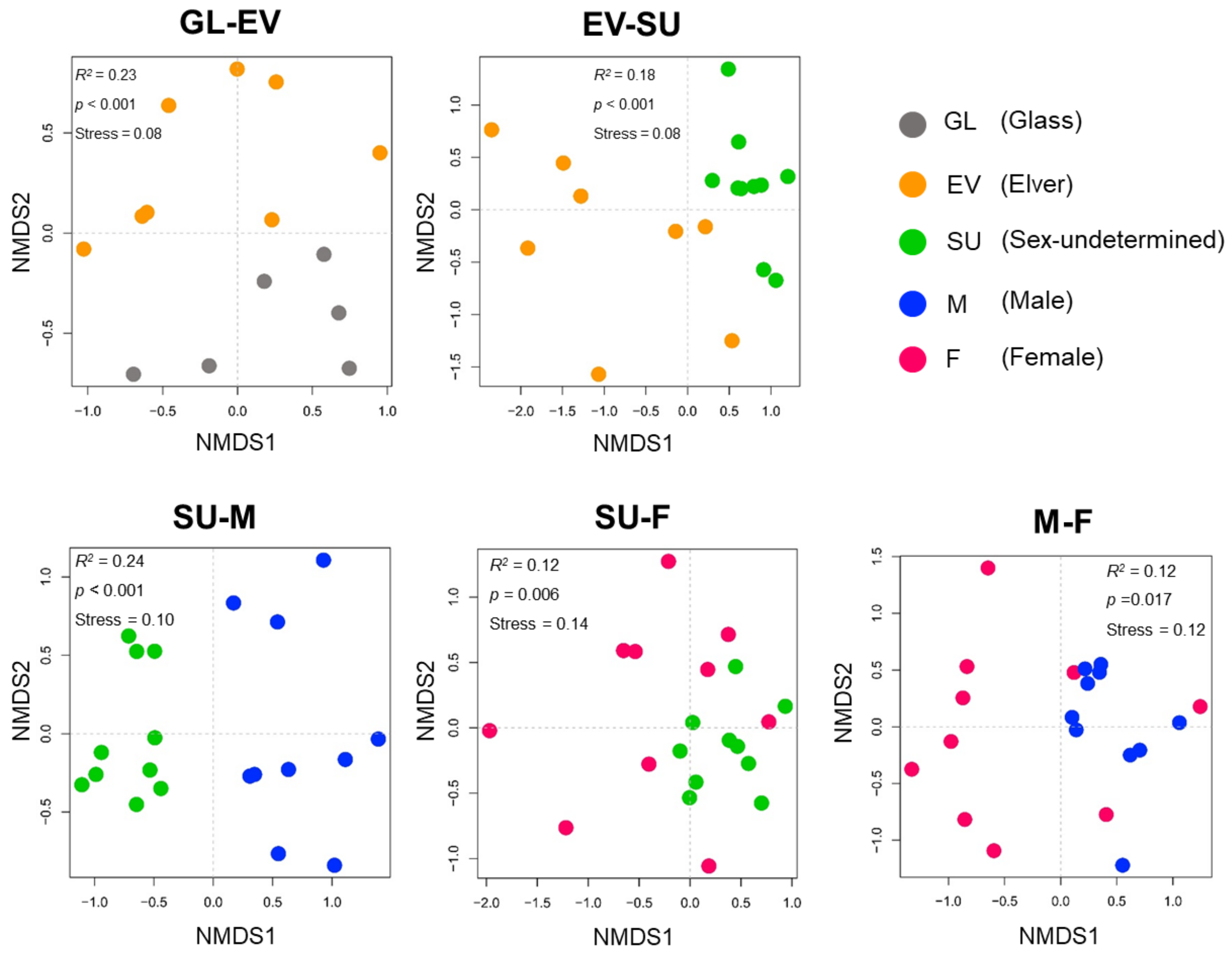


**Supplemental Figure 4**. Pairwise comparisons of gut miicrobial compositional structure between consecutive developmental stages (GL, EV, SU, M, and F) based on Bray-curtis dissimilarity and visualized in NMDS.

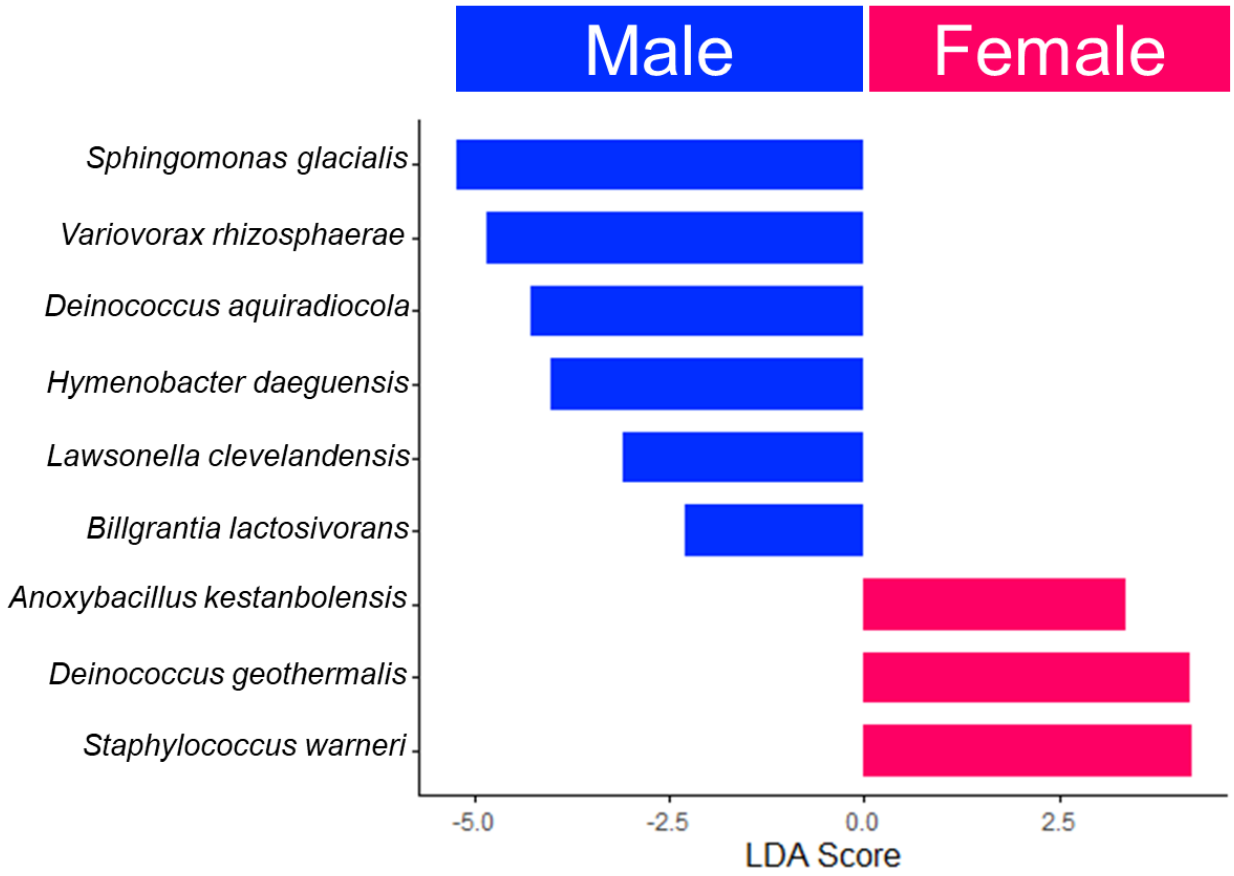


**Supplemental Figure 5.** LEfSe: Differential abundance of gut microbial genera between male and female eels.

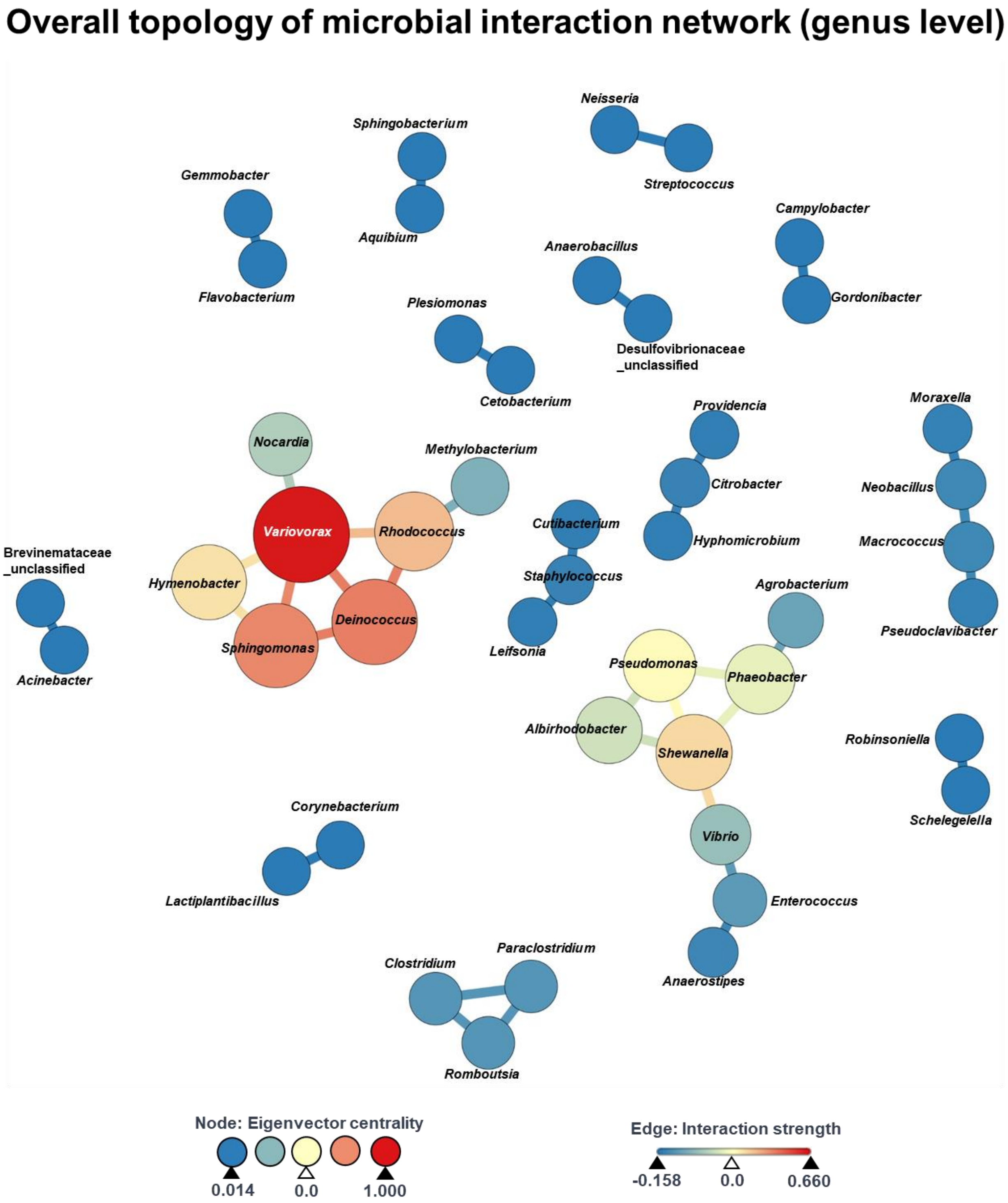


**Supplemental Figure 6.** Overall topology of microbial interaction network at the genus level across eel life stages. Each node represents a microbial genus, with node color and size indicating the eigenvector centrality score. Edge color represents the interaction strength between nodes, reflecting correlations between microbial genera. Correlations were calculated using FastSpar with 1000 bootstrap iterations.

**Supplemental Table 1.** Correlation analysis between bacterial genera across eel life stages. The top 10 bacterial species with the highest ranking based on eigenvector centrality score are shown in the table, together with degree centrality, closeness centrality, and betweenness centrality. Correlations were calculated using FastSpar with 1000 bootstrap replicates.

| Rank | Genus | Degree | Closeness | Betweenness | Eigenvector |
|---|---|---|---|---|---|
| 1 | *Variovorax* | 5 | 0.857 | 8.5 | 1.000 |
| 2 | *Deinococcus* | 3 | 0.667 | 1.0 | 0.782 |
| 3 | *Sphingomonas* | 3 | 0.600 | 0.5 | 0.761 |
| 4 | *Rhodococcus* | 3 | 0.667 | 5.0 | 0.649 |
| 5 | *Shewanella* | 4 | 0.636 | 13.0 | 0.588 |
| 6 | *Hymenobacter* | 2 | 0.546 | 0.0 | 0.569 |
| 7 | *Pseudomonas* | 3 | 0.500 | 1.0 | 0.514 |
| 8 | *Phaeobacter* | 3 | 0.500 | 6.0 | 0.455 |
| 9 | *Albirhodobacter* | 2 | 0.438 | 0.0 | 0.394 |
| 10 | *Nocardia* | 1 | 0.500 | 0.0 | 0.324 |

**Supplemental Table 2.** Correlation analysis between the abundance of bacterial genera and sex gene expressions (i.e., *amh* and *vtgr*).

| Gene expression | Genus | Rank | Correlation coefficient | P value |
|---|---|---|---|---|
| *amh* | *Sphingomonas* | 1 | 0.399 | 0.001 *** |
| | *Variovorax* | 2 | 0.359 | 0.002 * |
| | *Deinococcus* | 3 | 0.214 | 0.046 * |
| | *Romboutsia* | 4 | 0.167 | 0.087 |
| | *Clostridium* | 5 | 0.153 | 0.171 |
| | *Terrisporobacter* | 6 | 0.128 | 0.175 |
| | *Cutibacterium* | 7 | 0.109 | 0.286 |
| | *Hymenobacter* | 8 | 0.109 | 0.228 |
| | Tannerellaceae_Unclassified | 9 | 0.108 | 0.227 |
| | *Ezakiella* | 10 | 0.103 | 0.222 |
| *vtgr* | *Staphylococcus* | 1 | 0.214 | 0.056 |
| | *Cetobacterium* | 2 | 0.211 | 0.030 * |
| | *Macrococcus* | 3 | 0.161 | 0.085 |
| | *Aeribacillus* | 4 | 0.146 | 0.122 |
| | *Rhodococcus* | 5 | 0.135 | 0.184 |
| | Cyanobacteriota_Unclassified | 6 | 0.113 | 0.191 |
| | *Capnocytophaga* | 7 | 0.112 | 0.171 |
| | *Moraxella* | 8 | 0.100 | 0.264 |
| | Eubacteriales Family XII. Incertae Sedis_Unclassified | 9 | 0.094 | 0.272 |
| | *Abiotrophia* | 10 | 0.090 | 0.289 |
